# Foetal Bovine Serum as a Critical Hidden Variable in Neutrophil–Biomaterial Studies

**DOI:** 10.1101/2025.11.10.687474

**Authors:** Ezgi Irem Bektas, Jingzhi Fan, Gregor Miklosic, Vahid Jahed, Jacek K. Wychowaniec, Kristaps Kļaviņš, Matteo D’Este

**Author notes:** Corresponding author: Matteo D’Este, PhD, Address: Clavadelerstrasse 8, 7270 Davos Platz, Switzerland. **E-mail list: First author**: Ezgi Irem. **Co-authors:** Jingzhi Fan, Gregor Miklosic, Vahid Jahed, Jacek K. Wychowaniec, Kristaps Kļaviņš.

## Abstract

Immune regulation plays a crucial role during the regeneration process, and it determines the fate of inflammation after tissue injury or infection. Neutrophils serve as the primary immune cells recruited to the site of inflammation, initiating and directing the subsequent inflammatory cascade following implantation. As neutrophils are highly sensitive to environmental cues, commonly used culture supplements and molecules may themselves influence neutrophil function and consequently affect the interpretation of biomaterial-induced immune responses. This study examined how foetal bovine serum (FBS), a standard *in vitro* culture supplement, affects human peripheral blood neutrophils when supplied either in the culture medium or as a surface coating on 3D-printed polycaprolactone (PCL) scaffolds. The response to type I collagen coating was also assessed as a biologically relevant comparator. Neutrophil activity was evaluated by assessing metabolic activity and metabolomic profiles, reactive oxygen species (ROS) production, and inflammation-related markers via a high-throughput proximity extension assay. Type I collagen coating modified the metabolomic profile of neutrophils and MMP-9 release but had minimal effect on ROS generation. In contrast, the presence of FBS in the culture medium significantly influenced neutrophil behavior, leading to significant changes in metabolic activity, cytotoxicity, and the secretion of inflammation-associated molecules, even at concentrations as low as 1% (v/v). These findings highlight the importance of assessing the use of FBS in neutrophil culture models, particularly those isolated from humans, and emphasize the need to develop alternative platforms for investigating neutrophil–biomaterial interactions in a more physiologically relevant manner.

**Highlights:**

- Neutrophil response to foetal bovine serum (FBS) (1–10%) and collagen coatings on PCL scaffolds was tested.
- FBS impacts neutrophil activation and alters metabolite composition of the medium.
- FBS increased the release of inflammation-related molecules such as NE, IL-8 and VEGFA.
- Collagen changed neutrophil metabolites and decreased MMP-9 release.
- Results address the FBS bias and the need for physiologically relevant culture models.

**Graphical abstract:** (Created in BioRender.com (2025))

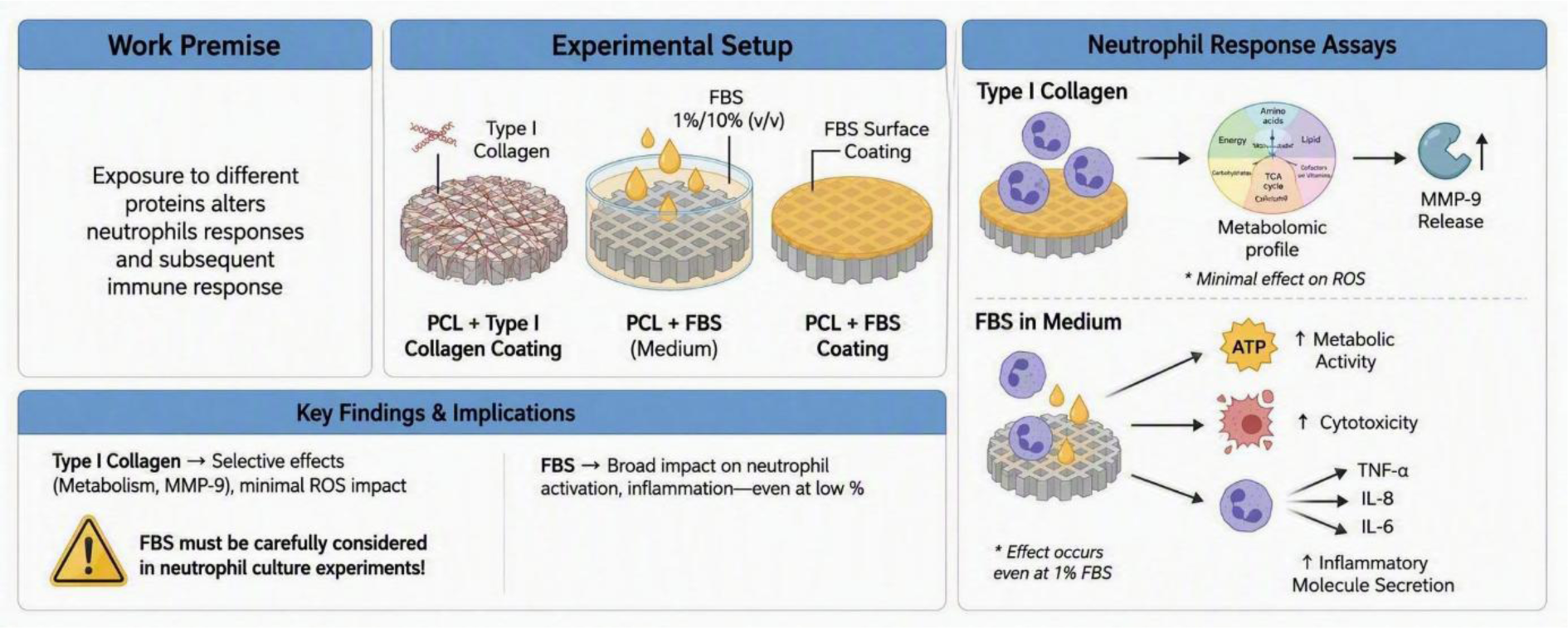

## 1. Introduction

Neutrophils are the most abundant type of leukocyte present in human peripheral blood, accounting for approximately 60% of the total white blood cell population [1]. They are primarily recognized as key initiators of acute inflammatory responses; additionally, neutrophils are pivotal in chronic inflammatory conditions, including inflammatory bowel disease, psoriasis, ulcerative colitis, atherosclerosis, and diabetes mellitus [2]. Due to their role as sentinels and first responders, neutrophils are critical mediators in determining the immunomodulatory properties of biomaterials through degranulation, apoptosis, neutrophil extracellular traps (NET) formation, cytokine release and recruitment of other immune cells [3]. Neutrophils are also instrumental in tumor pathogenesis, exhibiting phenotypic and functional heterogeneity across various cancer types, thereby influencing the tumor microenvironment in multiple ways. They can either induce tumor cell apoptosis by expressing tumor necrosis factor-related ligand and Fas ligand or facilitate malignant cell invasion by activating pathways such as JAK/STAT, which contribute to cancer cell proliferation, survival, invasion, and angiogenesis [4]. In the tumor microenvironment, neutrophils can adopt distinct phenotypes—N1 (antitumor) and N2 (protumor)—which are influenced by the molecular signals present in their surroundings. Beyond their roles in tumorigenesis, these phenotypes have also started to be investigated for their impact on tissue regeneration, wound healing, and the interplay between pro- and anti-inflammatory responses. Previous studies demonstrated that N1 neutrophils are characterized by increased production of reactive oxygen species (ROS), elevated levels of NADPH oxidase subunits, myeloperoxidase (MPO), and matrix metalloproteinase-9 (MMP-9), along with an enhanced chemotactic response, thereby confirming their classification as a pro-inflammatory phenotype [5]. By contrast, N2 neutrophils are classified as anti-inflammatory, polarizing macrophages toward an M2 phenotypes and promoting tissue regeneration [6]. Moreover, neutrophils have been shown to upregulate markers associated with osteogenesis—such as ALP, OCN, OPN, COL-I, and BMP2—as well as markers linked to angiogenesis, including VEGFA, VEGFB, and VEGFC, when co-cultured with human osteoblasts and human umbilical vein endothelial cells [7]. *Herath et al*. provided evidence supporting the role of neutrophils in enhancing bone tissue regeneration *in vivo* by repetitively administering autologous neutrophils to rabbits with calvarial defects [8]. Similarly, neutrophil-derived extracellular vesicles have been demonstrated to promote bone regeneration in both *in vitro* and *in vivo* models [9]. On the contrary, the presence of a high number of neutrophils due to hyper-inflammation decreased the matrix mineralization [10]. These findings indicate that, depending on the timing, severity of inflammation, and disease context, neutrophils become activated in different ways, undergo phenotypic changes (N1-N2), and regulate downstream events to facilitate tissue repair [11].

As the first responders to biomaterial implantation, neutrophils initiate and orchestrate the acute immune response by secreting cytokines and chemokines, recruiting macrophages and other immune cells, and attracting tissue-resident mesenchymal stromal cells (MSCs) to the site of inflammation [12]. Neutrophils are attracted by pathogen-associated molecular patterns (PAMPs) or migrate to the materials implantation site due to the presence of damage-associated molecular patterns (DAMPs), molecules released upon tissue damage, as well as the presence of blood proteins on the surface of the implanted materials [13]. The fate of inflammation is initially defined by the neutrophils and continues with the crosstalk between monocytes/macrophages and MSCs. Monocyte-derived macrophages differentiate into either an M1 (pro-inflammatory) or a subtype of M2 (anti-inflammatory) phenotype, depending on the microenvironment at the implantation site [14]. Macrophages are also involved in mediating the immunological response to biomaterials during tissue repair or regeneration [15]. Foreign body response (FBR) is one of the biological mechanisms in which the macrophages are involved via forming the foreign body giant cells (fusion of macrophages for phagocytosis of large objects) on the materials, triggered by the released molecules during the initial response and adsorption of serum proteins on the implants [16, 17]. When FBR becomes chronic, it promotes the formation of fibrous tissue and implant encapsulation [18]. Therefore, investigating how biomaterial properties modulate immune responses is crucial for developing design principles that account for their influence on the implant microenvironment.

The microenvironment plays a critical role in shaping neutrophil response, influencing their behavior, activation, and function. An *in vitro* model was previously introduced to characterize the neutrophil functions relevant to identifying their reaction to biomaterials [19]. Neutrophils are highly sensitive to various signals, including cytokines, growth factors, and extracellular matrix components, which guide their migration and activation. Therefore, the local microenvironment dictates the degree of neutrophil activation and determines their functions, including degranulation—where they release various proteins, cytokines, chemokines, and enzymes such as proteases (e.g., elastase, cathepsin) and myeloperoxidase—as well as the formation of neutrophil extracellular traps (NETs), phagocytosis, and efferocytosis [20, 21]. Hence, investigating the effects of different stimuli on neutrophils *in vitro* may further our understanding of how they orchestrate later stages of the immune response, contributing to deciphering their role in biomaterials-driven immunomodulation.

One important methodological consideration in this context is activation of neutrophils upon exposure to proteins, particularly serum proteins and collagen, which are ubiquitous in both biological systems and biomaterial-based applications. The use of foetal bovine serum (FBS) in neutrophil cultures remains controversial. Although FBS is a common supplement in cell culture systems [22], there is no agreement on the optimal concentration, with reported values ranging from 0.5 to 10% (v/v) [23–25]. Such variability in neutrophil-based *in vitro* assays may therefore introduce significant bias, especially in the growing number of studies evaluating the immunomodulatory properties of biomaterials, as neutrophils are highly responsive to serum-derived components, potentially leading to overactivation and misinterpretation of results. Type I collagen represents another biologically relevant stimulus because it is widely used in cell culture and tissue engineering applications and is among the most abundant extracellular matrix proteins encountered by immune cells during tissue repair. Despite their widespread use, the individual effects of serum proteins and collagen on neutrophil responses remain insufficiently understood.

To address this knowledge gap, we systematically investigated neutrophil responses to protein exposure associated with biomaterials by comparing the effects of FBS and type I collagen on polycaprolactone (PCL) surfaces. FBS was presented either as a supplement in the culture medium or as a coating on the biomaterial surface, thereby modelling both cell culture conditions and protein adsorption events that occur following implantation. Type I collagen was applied as a surface coating. PCL was selected as a model substrate because it is a widely used biodegradable polyester in tissue engineering, readily adsorbs proteins, and is suitable for 3D printing of relevant constructs. To elucidate changes in neutrophil function, we assessed metabolic activity, cytotoxicity, reactive oxygen species (ROS) production, inflammatory mediators, and intracellular metabolite profiles.

## 2. Materials and Methods

### 2.1 Fabrication of the 3D printed PCL and the preparation of the material coatings

Polycaprolactone (PCL) disc scaffolds were 3D printed (diameter = 10.6 mm) using pellets with a molecular weight (M_W_) of 45,000 Da (Sigma-Aldrich, Missouri, US) via an extrusion-based 3D printer, RegenHu 3D Discovery (RegenHU, Villaz-Saint-Pierre, Switzerland). The printing was performed with a nozzle of inner diameter 300 µm (G23 needle, RegenHu, Villaz-Saint-Pierre, Switzerland), and operated at 1 bar pressure, 25 revs/m rotation speed, and a writing speed of 6.5 mm/s. The temperature of the tank and printer head were set to 90°C and 80°C, respectively. After placing the scaffolds into the 48 well-plates, the discs were coated with either FBS (1% (v/v) or 10 % (v/v)) or type I collagen derived from rat tail (≈ 17 *μg*/scaffold) (354236, Corning, New York, USA). The discs were incubated overnight in FBS (35-079-CV, Corning, New York, USA) at +4 °C for coating. The type I collagen coating was performed according to manufacturer’s protocol. Briefly, the stock collagen was diluted with 0.02 N acetic acid solution to 50 *μ* g/mL concentration, dispersed and incubated onto disc scaffolds for 1h at room temperature (RT). Table 1 shows the experimental conditions to which neutrophils were subjected.

**Table 1.** The experimental conditions and their abbreviations.

| Abbreviations | Experimental groups |  |  |
| --- | --- | --- | --- |
|  | Material | Medium | Coating |
| **P** | PCL | RPMI 1640 | - |
| **P+F 1M** | PCL | RPMI 1640 + 1% (v/v) FBS | - |
| **P/F1** | PCL | RPMI 1640 | 1% (v/v) FBS |
| **P/F10** | PCL | RPMI 1640 | 10% (v/v) FBS |
| **P/C** | PCL | RPMI 1640 | Type I Collagen |
| **P/C+F 1M** | PCL | RPMI 1640 + 1% (v/v) FBS | Type I Collagen |
| **CP** | Cell culture plate | RPMI 1640 | - |
| **CP+F 1M** | Cell culture plate | RPMI 1640 + 1% (v/v) FBS | - |
| **CP+PMA** | Cell culture plate | RPMI 1640 + phorbol-12-myristate-13-acetate (positive control) | - |

The endotoxin levels of PCL discs were detected via Pierce™ Chromogenic Endotoxin Quant Kit (Thermo-Fisher, Waltham, Massachusetts, USA). The disc scaffolds were incubated in 1 mL of endotoxin free H_2_O (Sigma-Aldrich) for 1h at RT. The high (0.1-1.0 EU/mL) and low (0.01-0.1 EU/mL) standard E.coli endotoxin solutions were prepared and placed into the wells of the pre-warmed plate at 37°C. After transferring the samples to the plate wells, the Amebocyte Lysate Reagent was added onto the samples and incubated for the time indicated in the protocol. Following the addition of Chromogenic Substrate and the 6 minutes incubation time, the stop solution was added into all wells containing standards and samples for terminating the reaction. The optical density at 405 nm was measured using the Infinite® 200 Pro plate reader (Tecan, Männedorf, Switzerland).

### 2.2 Isolation, culture, and characterization of human peripheral blood neutrophils

Neutrophils were isolated from peripheral blood of 11 healthy human volunteers (aged 25-41 years) (Table S1) by immunomagnetic negative selection using the EasySep™ Direct Human Neutrophil Isolation Kit (STEMCELL Technologies, Vancouver, Canada) following the manufacturer’s instructions. The isolated cells were previously characterized and validated by flow cytometry analysis [19]. After the final step of the isolation procedure, following the centrifugation at 300 x g for 5 minutes, neutrophils were seeded with a density of 300,000 cells/scaffold, and cultured in Roswell Park Memorial Institute (RPMI) 1640 medium containing HEPES and L-Glutamine (Gibco, ThermoFisher, Waltham, USA), supplemented with 100 U/mL penicillin (Gibco) and 100 µg/mL streptomycin (Gibco).

### 2.3 Cell Titer Blue (CTB) metabolic activity assay

The metabolic activity of the neutrophils subjected to different types of scaffolds was measured using CellTiter-Blue® Cell Viability Assay (Promega, Dübendorf, Switzerland) which relies on the ability of living cells to convert the resazurin dye into the fluorescent compound resorufin. The CellTiter-Blue® Reagent was introduced into each well containing RPMI 1640 media at a ratio of 1:5 (v/v) 30 minutes before each respective time point (1h, 3h, 5h, 7h and 24h). The fluorescence intensity of the samples was recorded at 560_Ex_/590_Em_ using Infinite® 200 Pro plate reader (Tecan, Switzerland) after transferring reagent-media mixture to a 96 well plate.

### 2.4. Lactate dehydrogenase (LDH) Assay

The levels of LDH released into the cell culture supernatant due to the damage of the cellular plasma membrane were quantified using Cytotoxicity Detection Kit^PLUS^ LDH (Roche, Switzerland) following the manufacturer’s protocols. Briefly, the supernatants were taken at each respective time point and centrifuged at 300 g for 5 minutes at +4°C. The 100 µL from each sample was taken and transferred into the 96 well-plates. Then, the working solution (100 µL) was added into each well and incubated for 30 minutes at RT. After the addition of stop solution (50 µL), the absorbance values were determined at 490 nm via the Infinite® 200 Pro plate reader (Tecan, Switzerland). The cytotoxicity percentage was calculated for each sample applying the following formula: cytotoxicity (%) = 100 x (experimental value – low control)/(high control – low control) (Eq. 1)

The value for the low control was established using medium from cells that had been cultured on cell culture plates at the t=1h, and the high control value was determined using medium from cultured cells at each timepoint (1h, 3h, 5h and 24h), where 9% Triton X-100 was added 15 minutes before collecting the supernatants.

### 2.5 Measurement of superoxide anion (O_2_^•−^) and reactive oxygen species (ROS)

The amount of the superoxide anion produced by neutrophils was measured with the cytochrome c assay. The superoxide anions react with cytochrome c and are involved in the redox reaction wherein the ferricytochrome c (Fe^3+^) undergoes reduction, resulting in the formation of ferrocytochrome c (Fe^2+^). Briefly, 100 µM of cytochrome c (Cytochrome c from equine heart, C7752, Sigma-Aldrich) solution was prepared in Hanks’ Balanced Salt Solution (HBSS (no calcium, no magnesium, no phenol red), Gibco™, Waltham, USA) containing 10 mM HEPES (Gibco), 1 µM MgCl_2,_ 0.8 µM CaCl_2_, 100 U/mL penicillin (Gibco) and 100 µg/mL streptomycin (Gibco). The neutrophils were incubated in the prepared cytochrome c solution for 15 minutes and 4 hours (h). Following this, 200 µL of cell culture supernatant from each sample was transferred to a 96 well-plate, and then the absorbance values at 550 nm were measured and recorded. The concentration was calculated via Beer–Lambert’s law using the coefficient of 21.1 mM^-1^.cm^-1^. Superoxide dismutase (S5395, Sigma-Aldrich) at a concentration of 12.5 µg/mL was used as a control group to demonstrate the specificity of the assay for superoxide anions.

Luminometric analyses were performed to measure ROS production. Luminol enhanced chemiluminescence method enables the measurement of ROS, both intracellularly and extracellularly. The PCL scaffolds were 3D-printed with respect to the size of the white polypropylene 96 well-plates and placed into each well separately. After seeding the neutrophils (3×10^5^ cells/scaffold) into well-plates, 50 µM of luminol solution was prepared in RPMI 1640 media supplemented with 100 U/mL penicillin (Gibco) and 100 µg/mL streptomycin. FBS was also added with respect to the experimental groups in Table 1. The plate was placed into the pre-warmed (37 °C) Infinite® 200 Pro plate reader (Tecan, Switzerland) and the changes in the luminescence were recorded as relative light units (RLU) for 2 h at 3-minute intervals. The results were represented as the area under the curve (AUC).

### 2.6 Neutrophil Secretome Analysis

The neutrophil secretome was tested under the experimental conditions in Table 1. Neutrophil elastase and MMP9 protein levels were measured via human neutrophil elastase/ELA2 DuoSet and human MMP-9 DuoSet ELISA kits (R&D Systems, Minneapolis, USA) following the manufacturer’s guidelines on timepoints 1h, 5h and 24h. The absorbance values of each well were measured at 450 nm, and the wavelength correction was set to 540 nm.

The secretomes were also investigated for inflammation related human proteins with Olink® Inflammation reagent kit (Olink® Proteomics, Uppsala, Sweden) consisting of 92 proteins. The Olink method provides relative protein quantification with high specificity due to Proximity Extension Assay (PEA) technology in which 92 pairs of oligonucleotide-labelled antibody probes are designed to attach to their specific target proteins. The results were presented as Normalized Protein eXpression (NPX) values, which is an arbitrary unit in Log2 scale that is directly proportional to protein concentration. The data analysis was performed via online Olink Statistical Analysis App (https://olinkproteomics.shinyapps.io/OlinkStatisticalAnalysis/).

### 2.7 Metabolomics Analysis

The intracellular and extracellular metabolites of neutrophils were examined after incubation under the various conditions for 1, 5, and 24 hours. For extracellular samples, cell culture supernatants collected from each sample were centrifuged at 400 g for 5 minutes at 4 °C. Following centrifugation, the supernatants were mixed with pure methanol at a ratio of 1:4 (v/v) and vortexed. The intracellular samples were prepared by adding ice-cold methanol (80% (v/v)) on top of the cells and then scraping them for detachment. The samples were transferred into the Eppendorf tubes and centrifuged at 6000 rpm for 15 minutes at 4 °C. The final supernatants were collected, and both intra- and extracellular samples were stored at −70 °C.

For the analysis, the samples were dried using a vacuum concentrator (Vacufuge Plus, Eppendorf, Germany). The dried pellet was re-suspended with a solution mixing 90 µL of methanol (Supelco, LiChrosolv, USA) and 10 µL internal standard mixture (carnitine NSK-B-1 and amino acid MSK-A2-1.2, Cambridge Isotope Laboratories, USA), then transferred to the HPLC glass vials with inserts for further LC-MS analysis.

Targeted quantitative metabolite analysis was conducted using HILIC-based liquid chromatography and mass spectrometric detection. Metabolites were separated on analytical columns (ACQUITY Premiere BEH Z-HILIC 1.7 μm 2.1 x 100 mm, Waters) utilizing a gradient elution method with 0.15% formic acid and 10 mM ammonium formate in water as mobile phase A and a solution of 0.15% formic acid and 10 mM ammonium formate in 85% acetonitrile as mobile phase B with the total analysis time of 18 minutes. The mobile phase flow rate was set at 0.4 mL/min, injection volume at 2 μL, and column temperature at 40 °C. For MS detection, an Orbitrap Exploris 120 (ThermoFisher, Waltham, USA) mass spectrometer was used. The MS analysis was performed in ESI positive and ESI negative modes using full scan detection; the scan range was set from 50 to 600 m/z, and the mass resolution was set to 60000. The ESI spray voltage was set to 3.5 kV in positive mode and 2.5 kV in negative mode; the gas heater temperature was set to 400 °C; the capillary temperature was set to 350 °C; the auxiliary gas flow rate was set to 12 arbitrary units; and the nebulizing gas flow rate was set to 50 arbitrary units. For quantitative analysis, seven-point calibration curves with internal standardization were used. Tracefinder 5.1 General Quan (ThermoFisher, Waltham, USA) software was used for LC-MS data processing and quantification. Every reported metabolite was identified at level A [25] using an authentic standard compound previously mapped to the analytical system.

The metabolite data were analysed by MetaboAnalyst 6.0 and GraphPad Prism 10 [26]. The data were expressed as mean ± SD. The metabolite concentration data was normalized by Log transformation and Pareto scaling (mean-centered and divided by the square root of the standard deviation of each variable) for statistical analysis. Pathway analysis and enrichment analysis was performed using The Small Molecule Pathway Database as the pathway library.

### 2.8 Statistical Analysis

This study was carried out with a total of 11 donors (Table S1). Results are expressed as mean ± standard deviation. Statistical comparisons between groups were conducted using GraphPad Prism 10 (GraphPad Software Inc., USA). Metabolic activity, cytotoxicity, ROS production, and Olink® analyses were evaluated using a two-way ANOVA followed by Tukey’s multiple comparisons test, while ELISA results were assessed via one-way ANOVA with Tukey’s post-hoc test. Statistical significance is denoted in the figures as follows: *p < 0.05, **p < 0.01, ***p < 0.001, ****p < 0.0001.

## 3. Results & Discussion

Here we used a protocol of neutrophils isolation that was previously validated via flow cytometry showing 99.7% CD66b^+^ and 99% CD16^+^ cells, whereas the monocyte marker CD14 was detected on 0.2% [19]. The amount of endotoxin detected on PCL samples was 0.148 EU/mL (Table S2), which is below the FDA-recommended limit of 0.5 EU/mL for many medical devices.

### 3.1 FBS drives a metabolically activated neutrophil state with presentation mode-dependent changes in ROS production and secretome composition

Neutrophils are highly dynamic cells, responsive and sensitive to changes in their environment [21]. According to the CTB assay, neutrophils showed condition-dependent metabolic activity over the 24 h culture period (Figure 1a). At 1 h, neutrophils incubated on uncoated PCL showed higher metabolic activity than those on FBS-coated PCL, whereas adding FBS directly to the culture medium reduced metabolic activity at 1 and 3 h. Collagen-coated PCL did not differ markedly from uncoated PCL, and metabolic activity in neutrophils cultured on tissue culture plastic remained comparatively stable over 24 h. By 24 h, metabolic activity was lower across all experimental groups, reflecting the short lifespan of neutrophils, and the distinction between FBS-containing and FBS-free conditions diminished. Similarly, LDH analysis showed a significant increase across all groups at 24 h, indicative of cell membrane damage (Figure 1b). At 1, 3, and 5 h, FBS-containing groups generally released more LDH than FBS-free groups; however, at 24 h LDH release in FBS groups was lower except for CP+F 1M, where soluble FBS significantly increased LDH release. LDH results also indicated donor variability; for example, the higher “low control” (Eq. 1) values for donor 2 at 1 h of incubation led to cytotoxicity percentages below zero.

**Figure 1.**
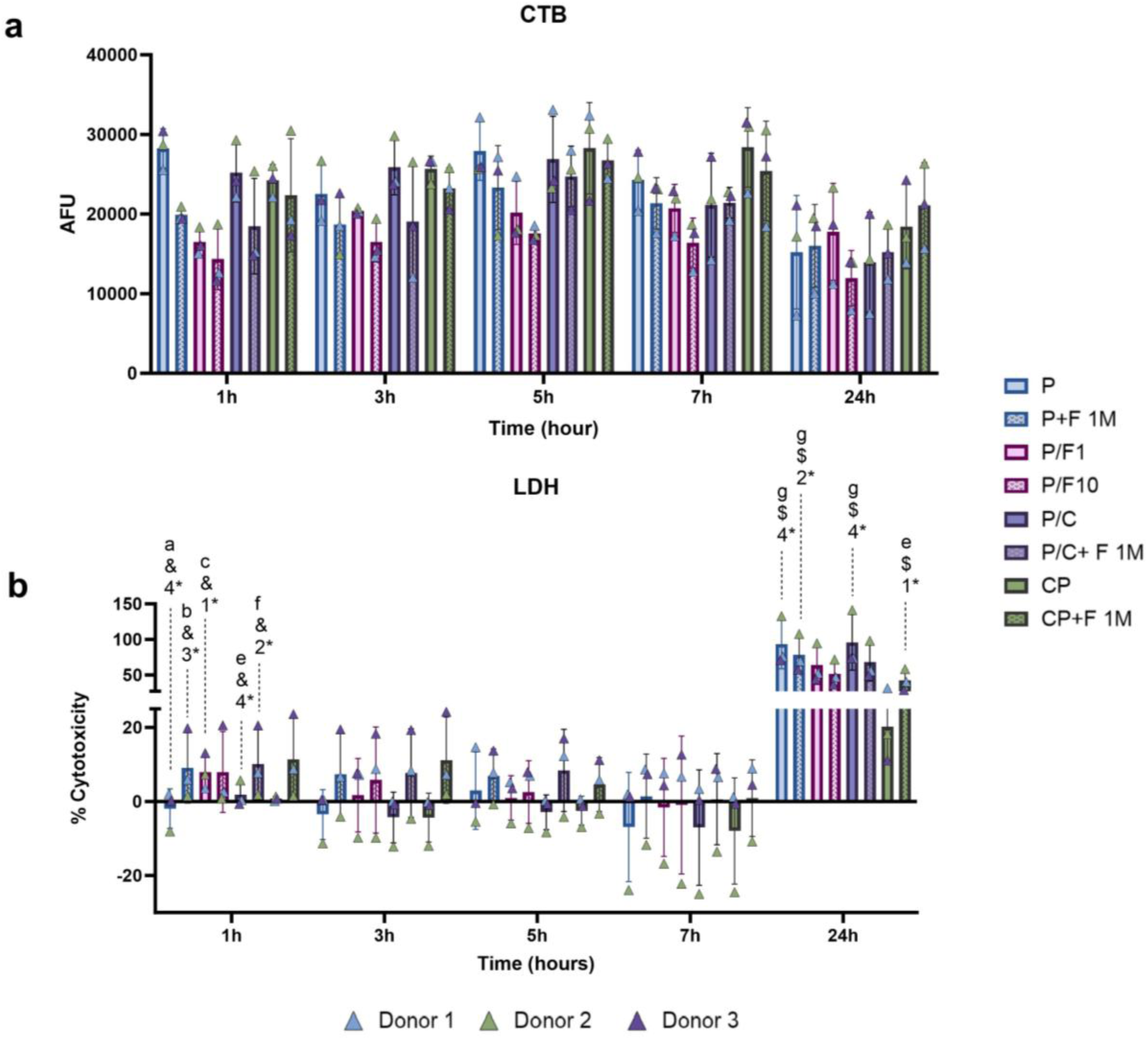
Neutrophil metabolic activity and LDH release on various substrates at 1h, 3h, 5h, 7h and 24h. a) Neutrophil metabolic activity assessed using the CellTiter-Blue assay, and the results were expressed as arbitrary fluorescence units (AFU) over time for each material. b) Neutrophils cultured on various substrates exhibited LDH release expressed as a percentage of the maximum LDH release following cell lysis. Data were obtained from three different biological donors (n=3). Column bars represent mean values +SD (SD: 1*: P< 0.05, 2*:P < 0.01, 3*:P < 0.001, 4*:P < 0.0001. Statistical significance is shown as; a: significantly different from the group P, b: significantly different from the group P+F 1M, c: significantly different from the group P/F1, e: significantly different from the group P/C, f: significantly different from the group P/C+F 1M, g: significantly different from the group CP, &: comparison of 1h-24h samples,$: comparison of 24h-24h samples).

We used cytochrome c and luminol enhanced chemiluminescence assays to measure extracellular superoxide anion and total intra- and extracellular ROS content [26], respectively. Because neutrophil activity peaked between 3 and 5 h, ROS measurements were performed at 4 h to represent the midpoint of this activation window. The specificity of the cytochrome c assay was confirmed by the absence of superoxide anions in the presence of superoxide dismutase (SOD), while the positive control, PMA, demonstrated increased ROS production as expected. Quantification at 4 h showed higher superoxide anion release under FBS-free conditions than in FBS-supplemented medium, and the net change between 15 min and 4 h further highlighted this temporal response (Figures 2a-b). Among the experimental groups, neutrophils cultured on P, P+F1M, and P/C showed the highest superoxide anion production, whereas FBS-coated PCL groups (P/F1 and P/F10) showed significantly reduced superoxide anion release. The mode of FBS administration also affected ROS production, with soluble FBS generally inducing higher levels than FBS used as a coating; conversely, adding FBS to collagen-coated cultures reduced superoxide anion production. Luminol-enhanced chemiluminescence revealed a similar trend for total ROS (Figures 2c-d), with FBS-coated PCL producing lower ROS than FBS-free groups, while collagen coating alone did not significantly alter ROS production. The elevated superoxide anion and total ROS production observed in the P and P/C groups may be attributed, at least in part, to the absence of FBS. Serum deprivation has previously been shown to induce ROS generation through ROS modulator 1 [27]. This mechanism may explain the increased ROS levels detected in these groups and their potential contribution to apoptosis [28]. However, the observation of similarly elevated ROS levels in the P + F 1M group indicates that FBS does not simply suppress oxidative stress but may exert a more complex and context-dependent influence on neutrophil activation.

**Figure 2.**
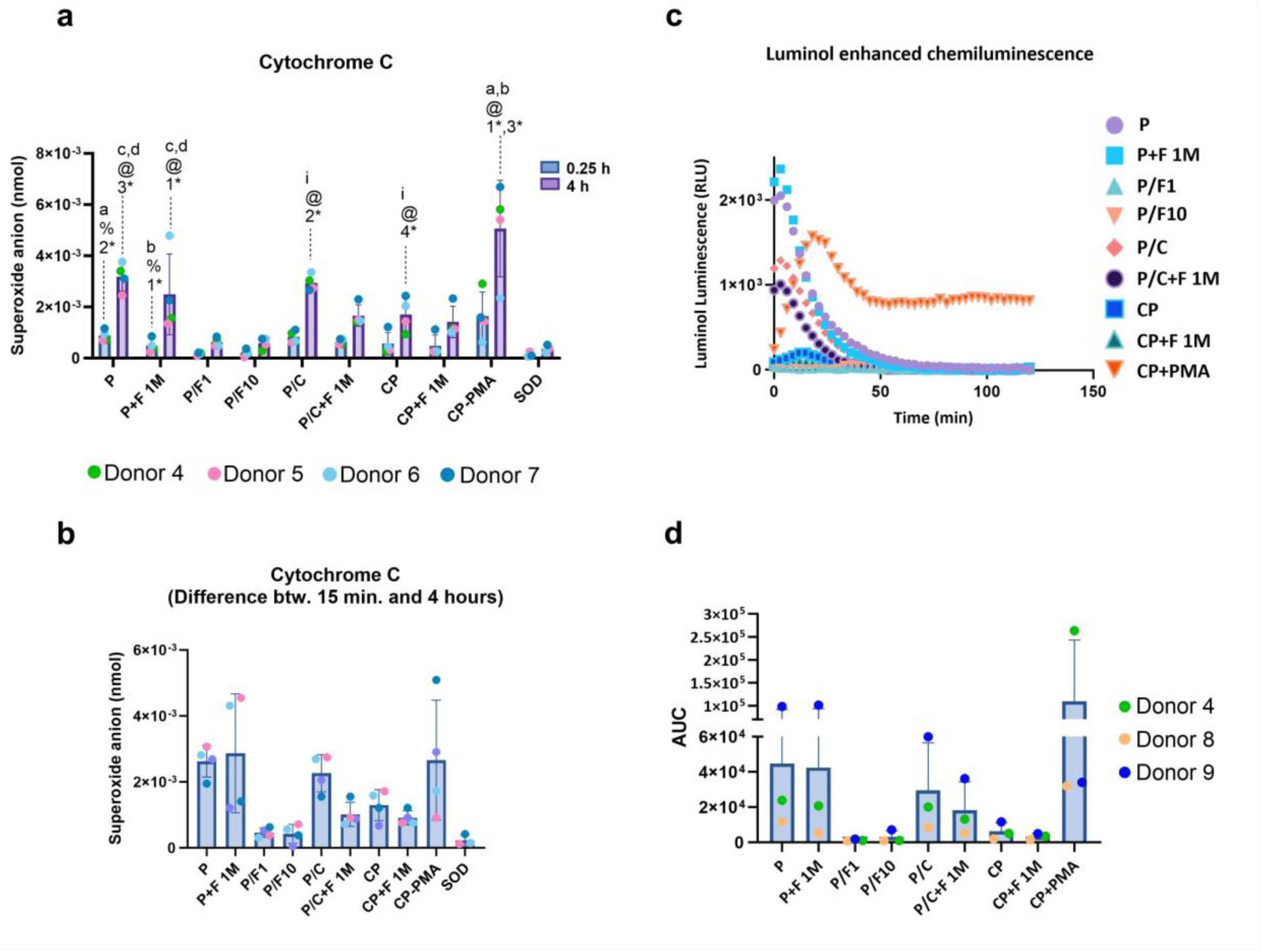
Effect of different biomaterials and coatings on ROS production in human peripheral blood neutrophils. (a-b) Superoxide anion was measured by cytochrome c reduction. (c) Total ROS production was measured through 2 hours and represented as relative light units (RLU) and, (d) the area under curve (AUC) was calculated from the RLU curves. SOD and PMA were used as negative and positive controls, respectively. Data were obtained from three different biological donors (n=3). Column bars represent mean values +SD (SD: 1*: P< 0.05, 2*:P < 0.01, 3*:P < 0.001, 4*:P < 0.0001. Statistical significance is shown as; a: significantly different from the group P, b: significantly different from the group P+F 1M, c: significantly different from the group P/F1, d: significantly different from the group P/F10, i: significantly different from the group CP+PMA %: comparison of 0.25h-4h samples, @: comparison of 4h-4h samples).

Analysis of the neutrophil secretome using the Olink® Inflammation panel revealed significant differential expression of 39 (Figure 3a) out of 92 inflammation-related proteins (Figure S1), primarily associated with innate immune processes, including neutrophil, monocyte, and leukocyte migration and chemotaxis (Figure 3b and Table 2). In this enrichment analysis, lower false discovery rate values indicated higher statistical significance, whereas higher signal values indicated a stronger association between the protein list and the corresponding biological process. CCL19 and CCL20 were consistently downregulated across all time points, while MCP-1, which is involved in migration and infiltration of monocytes and T cells [29], was upregulated at 5 and 24 h, particularly when FBS was present in the culture medium. As neutrophil recruitment is essential for coordinating innate and adaptive immune responses [30], alterations in chemotactic mediators may substantially influence downstream tissue responses.

**Figure 3.**
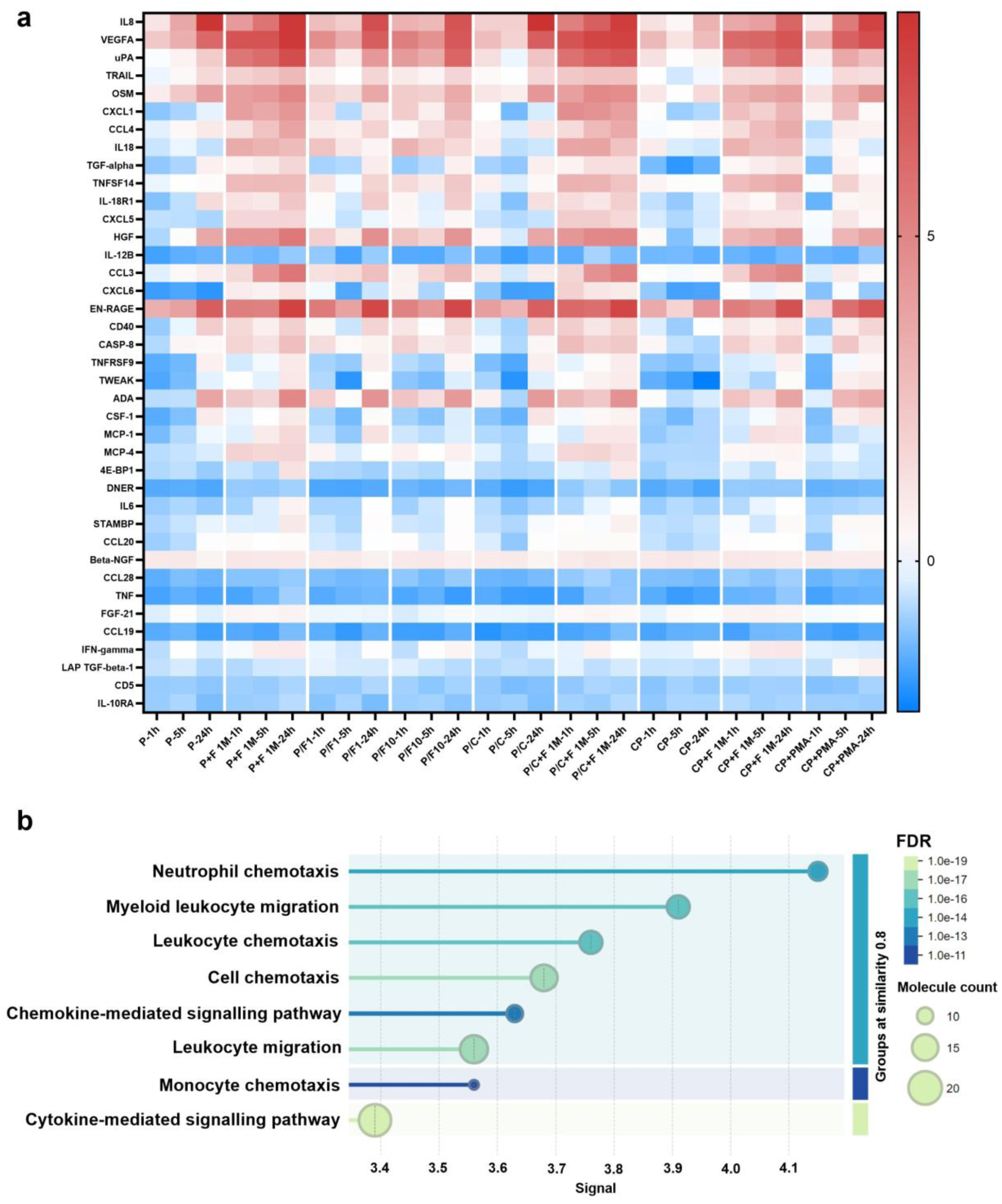
Comparison of the inflammation-associated proteins obtained from the neutrophil secretome of nine experimental groups through Olink analysis. a) Heatmap showing the expression of 39 different proteins from the Olink® inflammation panel quantified as NPX. Proteins included in the inflammation panel (rows) are displayed across the different experimental conditions and time points (1 h, 5 h, and 24 h; columns). Relative to external and plate controls, elevated protein levels are shown in red, whereas reduced levels are shown in blue. b) Biological process enrichment analysis of 39 significantly upregulated or downregulated proteins via https://string-db.org/, FDR: False discovery rate, Signal: weighted harmonic mean between the observed/expected ratio and −log (FDR). Data were obtained from three different biological donors (n=3) for each experimental group.

**Table 2.** Biological processes associated with proteins analyzed using Olink® platform (. https://string-db.org/).

| **Biological process** | **Proteins involved** |
| --- | --- |
| Neutrophil chemotaxis | EN-RAGE, CCL 3-4-19-20, MCP 1-4, CXCL1-5-6, IL-8 |
| Myeloid leukocyte migration | EN-RAGE, CSF1, MCP 1-4, CCL 3-4-19-20, CXCL1-5-6, IL-6, IL-8 |
| Leukocyte chemotaxis | TNFSF14, EN-RAGE, CCL 3-4-19-20, MCP 1-4, IL6, IL-8, CXCL1-5-6 |
| Cell chemotaxis | TNFSF14, EN-RAGE, HGF, CCL 3-4-19-20-28, MCP1-4, CXCL1-5-6, IL-6, IL-8 |
| Chemokine-mediated signalling pathway | MCP1-4, CCL3-4-19-20, CXCL1-5-6, IL-8 |
| Leukocyte migration | TNFSF14, EN-RAGE, CSF-1, CCL 3-4-19-20, MCP1-4, CXCL1-5-6, IL-6, IL-8, ADA, TNF |
| Monocyte chemotaxis | MCP 1-4, EN-RAGE, CCL3-4-19-20, IL-6 |
| Cytokine-mediated signalling pathway | CSF1, CCL 3-4-19-20, IL-6-8-18, OSM, CXCL1-5-6, MCP1-4, IL12B, TNF, IFNG, IL18-R1, IL10RA |

Among the most abundant proteins detected in the neutrophil secretome were IL-8, VEGFA, uPA, HGF, and EN-RAGE (Figure 4). IL-6 expression was downregulated in nearly all experimental groups, whereas IL-8 emerged as one of the most strongly affected proteins and was upregulated across groups, especially in the presence of FBS (Figure 4a). As a major neutrophil chemoattractant, IL-8 regulates neutrophil migration [31] and can modulate neutrophil phenotype in a concentration-dependent manner [6]. Therefore, the differences observed between FBS-coated and FBS-supplemented groups may contribute to distinct inflammatory and regenerative outcomes. Variability in IL-8 expression has also been reported between different FBS brands, raising concerns regarding experimental reproducibility in immune-related studies [22]. The increased IL-8 levels may have stimulated NE release in FBS-treated groups [32, 33] and, via its angiogenic activity [34], contributed to the increase in VEGFA (Figure 4b) [35–37]. FBS exposure also markedly increased the secretion of uPA (Urokinase-type Plasminogen Activator, involved in the conversion of plasminogen into plasmin, Figure 4c) and HGF (Figure 4d), particularly at 1 and 5 h when FBS was supplied in the culture medium rather than as a coating. At 24 h, uPA maintained this trend, whereas VEGFA and HGF (Vascular Endothelial Growth Factor A and Hepatocyte Growth Factor, respectively) differences were less distinct. EN-RAGE levels remained largely unchanged between groups, although a slight increase was detected in the presence of soluble FBS (Figure 4e). Through its interaction with uPAR (uPA receptor), uPA initiates plasmin-mediated degradation of fibrin and extracellular matrix components [38]. Beyond its role in tissue remodeling, uPA has been implicated in cartilage damage in rheumatoid arthritis [39] and impaired neutrophil efferocytosis by macrophages [40], suggesting a potential contribution to the persistence of inflammation. In contrast, HGF is associated with cell proliferation, growth and migration [41], and has been shown to regulate MSC function [42]. Neutrophil-derived HGF has been linked to regenerative responses following tissue injury [43], while HGF-MET signaling promotes macrophage polarization toward the pro-regenerative M2 phenotype [44]. Thus, increased HGF release may represent a compensatory mechanism favoring tissue repair. EN-RAGE, although not significantly altered by FBS treatment, remains relevant due to its established role in chronic inflammatory diseases, including rheumatoid arthritis, inflammatory bowel disease, diabetes, and cancer [45]. As both a ligand and regulator of RAGE signaling, EN-RAGE can inhibit neutrophil migration [46] and impair function [47], and has been implicated in elevated NET formation [46], highlighting its potential contribution to neutrophil-mediated inflammatory responses. Overall, neutrophils cultured in the presence of FBS secreted more inflammation-related proteins than those cultured without FBS, and soluble FBS generally induced a stronger protein release response than FBS presented as a coating, especially at early time points.

**Figure 4.**
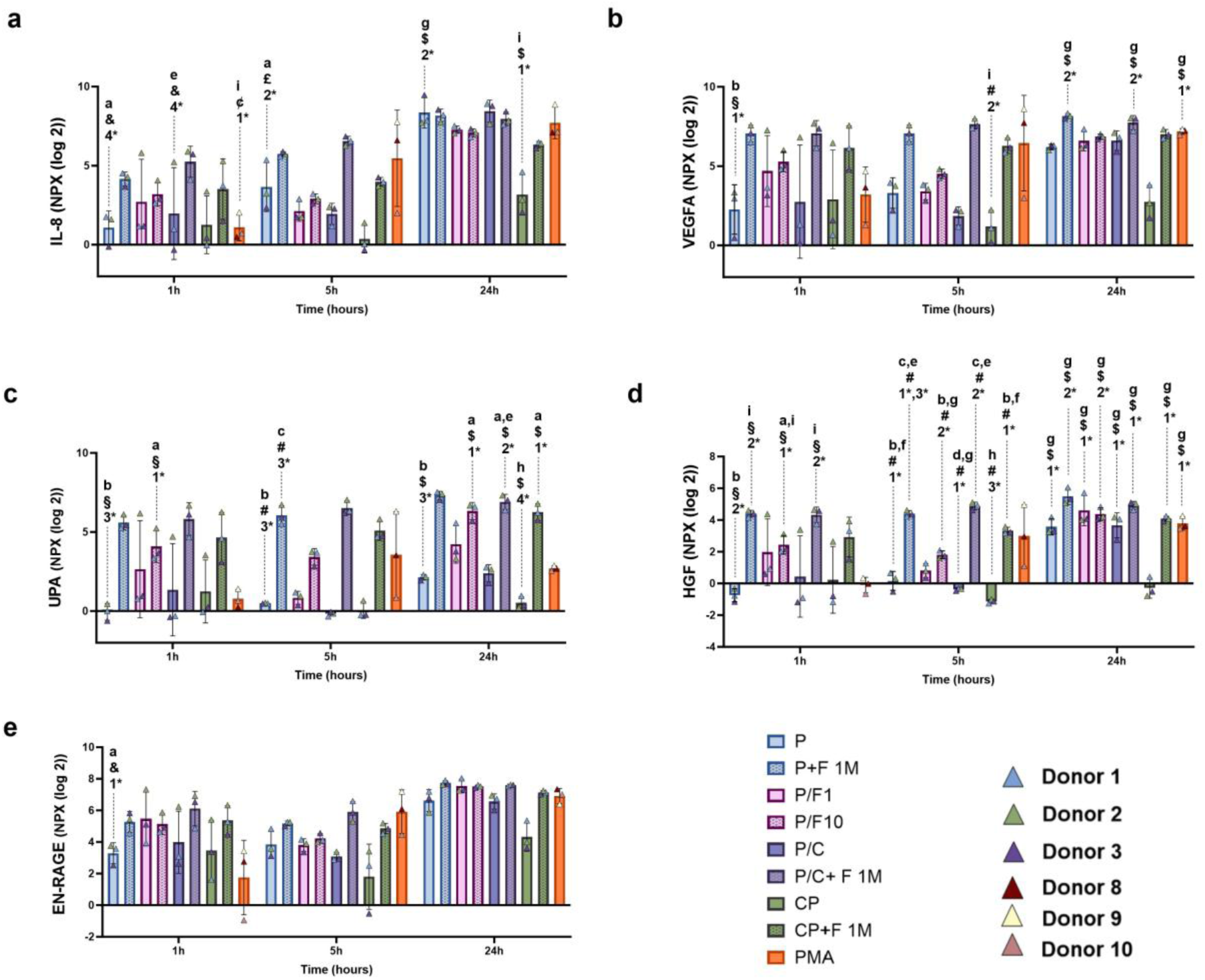
NPX (log2) column plots of a) IL-8, b) VEGFA, c) UPA, d) HGF, e) EN-RAGE. Data were obtained from three different biological donors (n=3) for each experimental group. Column bars represent mean values ±SD (SD: 1*: P< 0.05, 2*:P < 0.01, 3*:P < 0.001, 4*:P < 0.0001). Statistical significance is shown as; a: significantly different from the group P, b: significantly different from the group P+F 1M, c: significantly different from the group P/F1, d: significantly different from the group P/F10, e: significantly different from the group P/C, f: significantly different from the group P/C+F 1M, g: significantly different from the group CP, h: significantly different from the group CP+F 1M, i: significantly different from the group CP+PMA. (**§:** comparison of 1h-1h samples, **#:** comparison of 5h-5h samples, **$:** comparison of 24h-24h samples, **&:** comparison of 1h-24h samples, **¢:** comparison of 1h-5h samples, **£:** comparison of 5h-24h samples).

MMP-9, a neutrophil-derived protease involved in inflammation, tissue remodeling, angiogenesis, and chemotaxis [48], was elevated in all FBS-exposed groups and in group P compared with P/C and CP (Figure 5a). Collagen coating significantly reduced MMP-9 release from PCL scaffolds, whereas soluble FBS produced a more pronounced increase in collagen-coated PCL and tissue culture plastic groups. ROS may partly drive this response in groups P, P + F 1M, and P/C [49], while the remaining increases are more consistent with direct FBS-induced secretion. FBS has been shown to enhance MMP-9 expression in foetal membrane cells [50], potentially through increased synthesis of prostaglandin, an inflammatory mediator that can also be produced by activated neutrophils [51]. ROS are also known to promote NETosis by facilitating granule permeabilization and the translocation of NE to the nucleus, where it promotes chromatin decondensation [52, 53]. NE release was elevated in FBS-containing groups, consistent with the patterns observed for ROS and MMP-9 (Figure 5b). Although collagen displayed a similar decreasing trend for NE as observed for MMP-9, this effect was not statistically significant. However, the reduced NE levels detected in group P, despite enhanced ROS production, indicate that oxidative stress under serum-free conditions may not translate into activation of NETosis. Instead, ROS may preferentially drive alternative cell death pathways [54]. Furthermore, the method of FBS presentation appeared to influence neutrophil behavior, as surface-coated FBS induced slightly lower NE release than FBS supplied directly in the culture medium. Given that extracellular NE release from azurophilic granules is a marker of neutrophil activation and inflammation [55], these findings indicate that soluble FBS components may stimulate a stronger inflammatory response than coated proteins. Collectively, these findings suggest that both oxidative stress and FBS-dependent signaling contribute to neutrophil activation, although their relative contributions may vary depending on scaffold properties and the mode of protein presentation.

**Figure 5.**
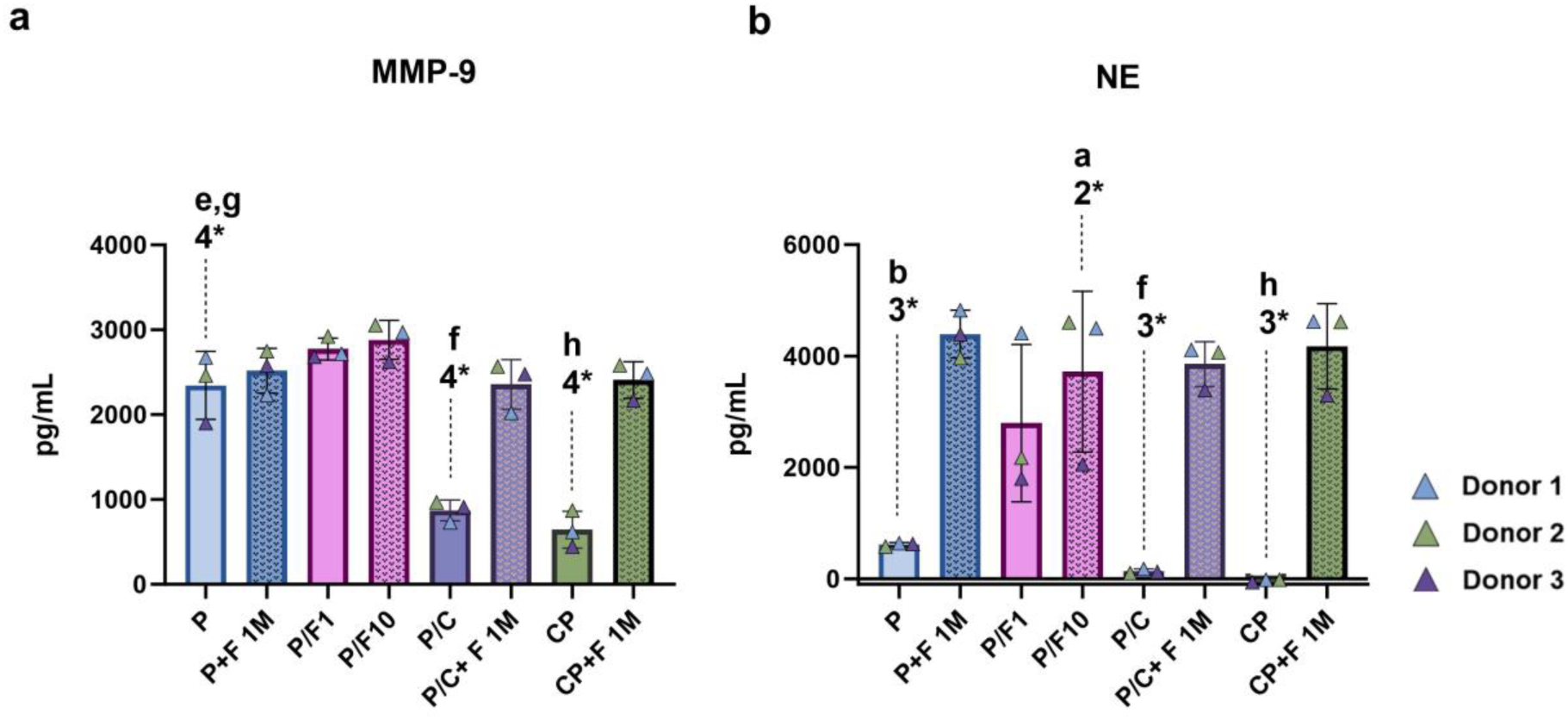
Quantification of a) MMP-9 and b) NE in the cell culture medium under various experimental conditions. Data were obtained from three different biological donors (n=3) for each experimental group. Column bars represent mean values ±SD (SD: 2*:P < 0.01, 3*:P < 0.001, 4*:P < 0.0001). Statistical significance is shown as; a: significantly different from the group P, b: significantly different from the group P+F 1M, e: significantly different from the group P/C, f: significantly different from the group P/C+F 1M, g: significantly different from the group CP, h: significantly different from the group CP+F 1M.

Intracellular metabolomic profiling performed at 1, 5 (Figure 6), and 24 h (Figure S2) revealed marked metabolic alterations in neutrophils exposed to FBS, either as a coating or in the culture medium. Principal component analysis further showed distinct clustering between FBS-free groups (P, P/C, and CP) and groups exposed to soluble FBS at 1 h (Figure S3a), while collagen- and FBS-containing groups remained separated from untreated controls at 5 h (Figure S3b). At 24 h, increased overlap among several groups suggested metabolic convergence associated with cellular exhaustion or death (Figure S3c). The distinct metabolomic signatures observed across experimental groups indicated that neutrophil response is changed in a time-dependent manner, as further supported by the pathway enrichment analysis conducted on the neutrophil samples (Figures S4-7). Groups CP+F1M (Figures 6a-b) and P+F1M (Figures 6c-d) exhibited reduced levels of energy metabolites (ATP, ADP, and AMP) and TCA cycle intermediates (citric acid, isocitric acid, and α-ketoglutaric acid), suggesting a shift from oxidative metabolism toward glycolysis. The early depletion of adenylates likely reflects the acute functional demand of activated neutrophils, whereas by 5 h increased lactate levels and partial recovery of ATP/ADP with suppressed AMP suggest glycolytic compensation and nucleotide salvage that restore adenylate energy charge (Figure S8). This metabolic reprogramming may also explain the reduced CTB signal observed in FBS-treated groups (Figure 1a), as the assay relies on NADH/NADPH-dependent reduction of resazurin. Increased consumption of NADPH during the respiratory burst may further contribute to the elevated ROS production, particularly in the P+F1M group (Figure 2). Interestingly, neutrophils cultured on FBS-coated PCL scaffolds displayed a pronounced reduction in taurine levels at 1 h, which was not observed in other FBS-containing groups (Figures 6a and e). Taurine is converted by activated neutrophils into taurine chloramine (Tau-Cl), a molecule that suppresses MPO degranulation and limits ROS-dependent tissue damage and NET formation [56, 57]. Taurine has also been reported to inhibit superoxide and nitric oxide generation and reduce pro-inflammatory cytokine production [58, 59], which may explain the lower superoxide anion and total ROS levels observed in neutrophils cultured on FBS-coated scaffolds (Figure 2). FBS-treated groups also showed consistently elevated levels of creatinine, butyrylcarnitine, and uridine (Figures 6c-f). Increased creatinine, a by-product of phosphocreatine metabolism, may reflect enhanced energy demand [60] associated with neutrophil activation and degranulation, consistent with the increased MMP-9 and NE release observed in these groups (Figure 5). Similarly, elevated butyrylcarnitine levels suggest altered fatty acid metabolism [61, 62]. Although neutrophils predominantly rely on glycolysis, fatty acid oxidation contributes to functions such as degranulation, phagocytosis, chemotaxis, and NET formation [63]. Accumulation of butyrylcarnitine may therefore indicate increased fatty acid utilization or incomplete β-oxidation during activation. Uridine was likewise elevated following FBS exposure. Its accumulation may reflect increased nucleotide turnover during neutrophil activation. Although extracellular uridine nucleotides promote neutrophil recruitment through P2Y₂ receptor signaling [64], uridine itself can suppress neutrophil infiltration, pro-inflammatory cytokine expression, and ROS production [65]. Thus, increased uridine may reflect an endogenous regulatory response that counterbalances FBS-induced neutrophil activation. Taken together, the metabolomic alterations observed in FBS-treated groups are consistent with the enhanced inflammatory and secretory responses identified throughout this study. The depletion of energy metabolites, accumulation of activation-associated metabolites, and changes in taurine metabolism suggest that neutrophil metabolic reprogramming is closely linked to ROS production, degranulation, and cytokine release, highlighting the importance of considering serum-derived effects when evaluating neutrophil–biomaterial interactions.

**Figure 6.**
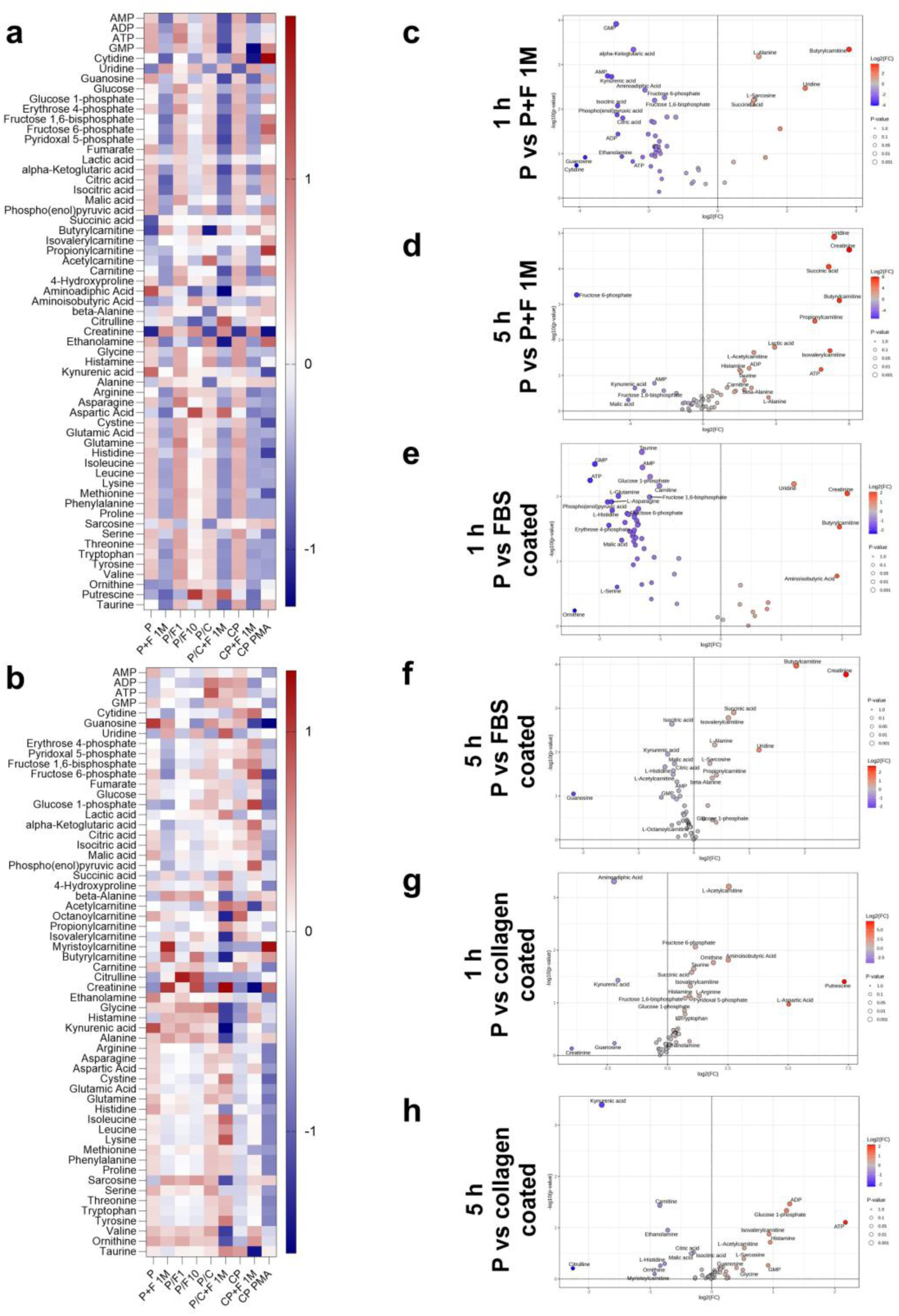
**Intracellular** metabolite profiles of neutrophils cultured under different in vitro conditions at defined time points. Heatmaps representing the upregulated (red) and downregulated (blue) metabolites (in rows) in neutrophils after a) 1 hour, b) 5 hours of incubation under various experimental conditions (in columns) and volcano plots illustrating the fold changes (log₂FC) versus statistical significance (–log₁₀p-value) of detected metabolites in neutrophils under different conditions at c) 1 hour P vs P+F 1M, d) 5 hours P vs P+F 1M, e) 1 hour P vs FBS coated P, f) 5 hours P vs FBS coated P, g) 1 hour P vs collagen coated P, h) 5 hours P vs collagen coated P.

### 3.2 Type I Collagen Coating Modulates Neutrophil Metabolism and Reduces MMP-9 Release with Minimal Effects on ROS Generation

Type I collagen alone maintained neutrophil metabolic activity, indicating that the coating itself was well tolerated by the cells (Figures 1a-b). While collagen alone did not compromise neutrophil viability, it modulated neutrophil function, resulting in elevated superoxide anion production. This response may be mediated by collagen-induced activation of β2 integrins, which has previously been linked to superoxide anion generation in neutrophils [66, 67]. In addition, exogenous collagen can promote platelet aggregation and subsequent production of superoxide anion and hydroxyl radicals [68]. As collagen-induced ROS further enhance platelet activation [69], these mechanisms may overall contribute to the increased oxidative response observed in collagen-coated conditions.

Collagen exposure induced rapid metabolic reprogramming in neutrophils during the early hours (1 and 5 h), characterized by enhanced glycolysis, glycogenolysis, and ROS production (Figures 6a, b, g and h) [70, 71]. In the first hour of collagen stimulation, neutrophils exhibited elevated glycolytic intermediates, including fructose 6-phosphate, fructose 1,6-bisphosphate, and glucose 1-phosphate, together with increased succinate, carnitine derivatives, and urea cycle–associated metabolites such as ornithine and putrescine (Figures 6a and g). This metabolic phenotype was further supported by the accumulation of succinate, carnitine derivatives, and metabolites associated with arginine and polyamine metabolism, which may be due to increased utilization of multiple energy sources to sustain inflammatory functions and NET formation [72–75]. However, this early metabolic activation was transient, as neutrophils cultured on collagen-coated scaffolds shifted from a highly active effector phase at 5 h (Figures 6B and H) to marked depletion of energy metabolites by 24 h (Figures S9a and d), indicating metabolic exhaustion following sustained activation.

Uncoated PCL induced early increases in kynurenic acid, guanosine, creatinine, aminoadipic acid, and uridine, alongside reductions in fructose-6-phosphate and related metabolites (Figure S9a-c). The elevation of kynurenic acid and guanosine, together with reduced NE release (Figure 5), suggests a counter-regulatory response that limits neutrophil activation, degranulation and NETosis [76–78]. In contrast, PMA-treated neutrophils exhibited a progressive metabolic shift characterized by transient accumulation of glycolytic intermediates followed by depletion of energy-related metabolites, consistent with sustained activation and eventual metabolic exhaustion (Figure S10). PCL degradation may also contribute to neutrophil metabolism, as hydrolysis produces 6-hydroxyhexanoic acid [79], which can be converted to acetyl-CoA (Figure S11a) that promotes a shift toward TCA cycle activity, as reflected by increased citric acid and isocitric acid at 24 h (Figures S7c and S9a). In parallel, declining lactic acid levels from early exposure to 5 and 24 h (Figure S11b) suggest a shift from glycolysis toward oxidative metabolism. Decreases in ATP, ADP, glucose 1-phosphate, and carnitine at 5 h incubation (Figure S9b) indicate high energy consumption, whereas increased amino acids may support antioxidant defense and controlled inflammatory responses. Elevated α-aminoadipic acid (Figure S9c) further indicates oxidative protein damage largely through neutrophil MPO activity [80].

The overall response of neutrophils incubated on PCL scaffolds and with type I collagen coatings was less pronounced than that of neutrophils exposed to FBS, whether as a coating or within the cell culture medium. The only observed difference was a slightly higher ROS production in neutrophils seeded on PCL scaffolds (groups P, P+F1M, P/C) compared to those on tissue culture plastic (groups CP and CP+F 1M).

### 3.3. Study limitations

One limitation of our study was the use of serum without heat inactivation(i.e., holding the serum at 56 °C for 30 minutes). The choice was to limit the loss of biological activity of components such as amino acids, vitamins, and growth factors [81, 82]. Previous research has demonstrated that heat inactivation of serum causes protein aggregation [83], changes in protein corona composition and cellular uptake of nanoparticles [84, 85]. Linscott *et al.* investigated the bovine complement system and revealed that heat inactivation affects various complement components differently. Specifically, they found that while complements C1, C2, and C8 are heat sensitive, leading to their inactivation at 56°C, the levels of C3 and C6 increased post heating. This rise is attributed to the inactivation of heat-labile inhibitors that normally regulate these components [86]. In another study, it was reported that heat-stable nucleases present in serum remained active even after heat inactivation at 56°C, leading to the degradation of NETs [87]. In summary, while heat-inactivated serum is a common practice in immunological experiments to neutralize complement proteins, aforementioned studies indicate that this practice could modify other FBS components, potentially affecting neutrophil responses and leading to data misinterpretation. Another notable limitation is that immune regulation comprises a highly complex network of biochemical pathways and multicellular interactions that cannot be fully reproduced in vitro. Nevertheless, reductionist approaches remain valuable for elucidating key mechanisms and isolating specific factors that contribute to the overall response.

## 4. Conclusions

In this study, we evaluated the response of human neutrophils to FBS and type I collagen coatings, as well as to FBS supplementation in the culture medium. To our knowledge, this is the first study to provide a detailed characterization of FBS effects on neutrophils in the context of widely used 3D-printable and implantable PCL biomaterials. Our results demonstrate that the presence of FBS, whether applied as a coating or added to the medium, markedly altered neutrophil behavior, affecting metabolic activity, cell membrane integrity, and the secretion of inflammation-related molecules, even at the lowest concentration tested (1 % (v/v)). FBS exposure also induced significant shifts in metabolite levels, including increases in creatinine, butyrylcarnitine, and uridine during the early time points (1–5 h). In contrast, type I collagen primarily modified the neutrophil metabolomic profile, showing elevated glycolytic intermediates along with increased succinate, carnitine derivatives, and urea cycle–associated metabolites such as ornithine and putrescine, while exerting minimal influence on ROS production in comparison to the response observed with PCL alone. Collectively, these findings underscore the importance of carefully evaluating protein presentation to neutrophils *in vitro*, as it can profoundly influence experimental outcomes. Building on this foundation, future investigations will aim to elucidate how the neutrophil secretome orchestrates cell recruitment and tissue regeneration. These insights lay the groundwork for advancing our understanding of biomaterial-mediated immunomodulation.

## Supporting information

Supplementary material Bektas et al.

