## Supplementary material Bektas et al. for "Foetal Bovine Serum as a Critical Hidden Variable in Neutrophil–Biomaterial Studies"

**Table. S1** Summary of donor information for experimental analyses.

| **Experiments** | **Donors** | **Sex** |
| --- | --- | --- |
| CTB, LDH, MMP-9, NE | 1,2,3 | Female |
| Cytochrome C | 4,5,6,7 | 2 females, 2 males |
| Luminol enhanced chemiluminescence | 4,8,9 | 1 female. 2 males |
| Olink® | 1,2,3-8,9,10 | 5 females, 1 male |
| Metabolomics analysis | 11 | 1 female |

**Table. S2** The amount of endotoxin detected on PCL samples.

| | PCL Sample 1 | PCL Sample 2 | PCL Sample 3 | |
| --- | --- | --- | --- | --- |
| Endotoxin concentration (EU/mL) | 0.143 | 0.145 | 0.157 | Average: 0.148 |

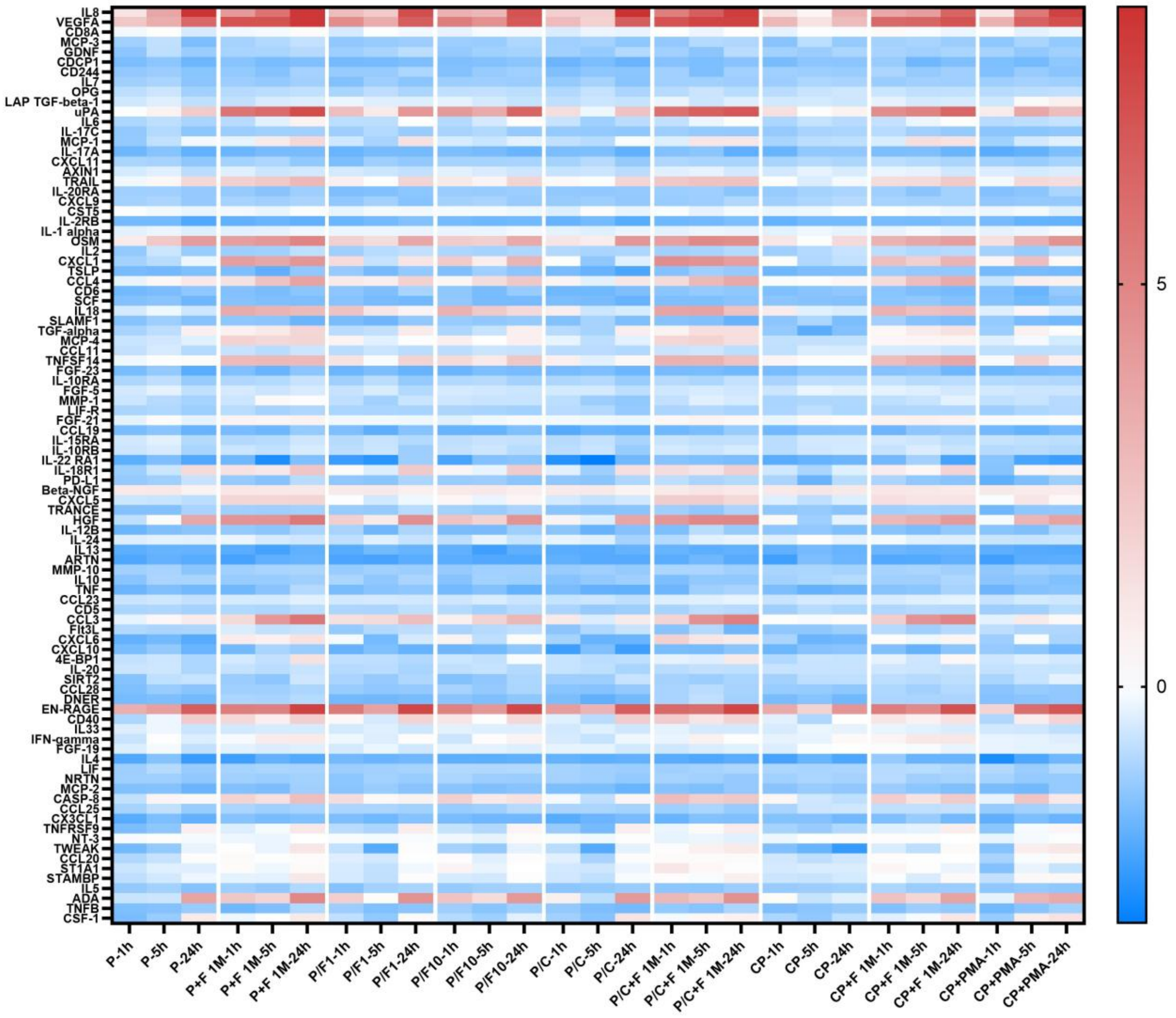


**Figure S1.** Heatmap showing the expression of 92 different proteins from the Olink® inflammation panel quantified as normalized protein expression (NPX; (log2 scale)). Proteins included in the inflammation panel (rows) are displayed across the different experimental conditions and time points (1 h, 5 h, and 24 h; columns). Red indicates higher, whereas blue represents lower protein levels relative to external and plate controls.

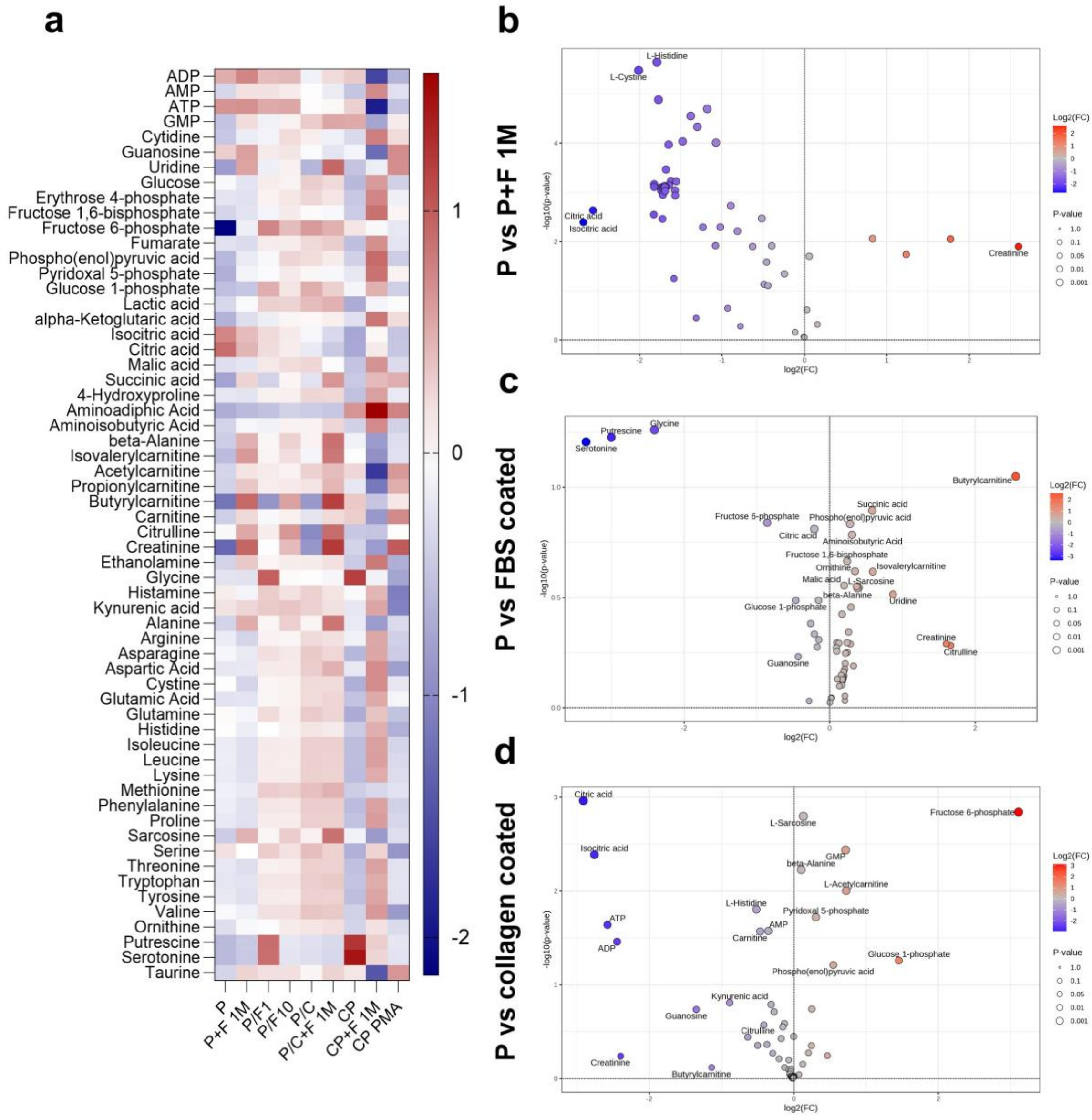


**Figure S2.** Metabolomic profiles of neutrophils cultured under different in vitro conditions at 24 hours. a) Heatmap representing the significantly upregulated and downregulated metabolites in neutrophils after 24 hours. Volcano plots showing differential metabolite expression on 24[th] hour of the comparisons b) P vs P+F 1M, c) P vs FBS coated, d) P vs collagen coated P.

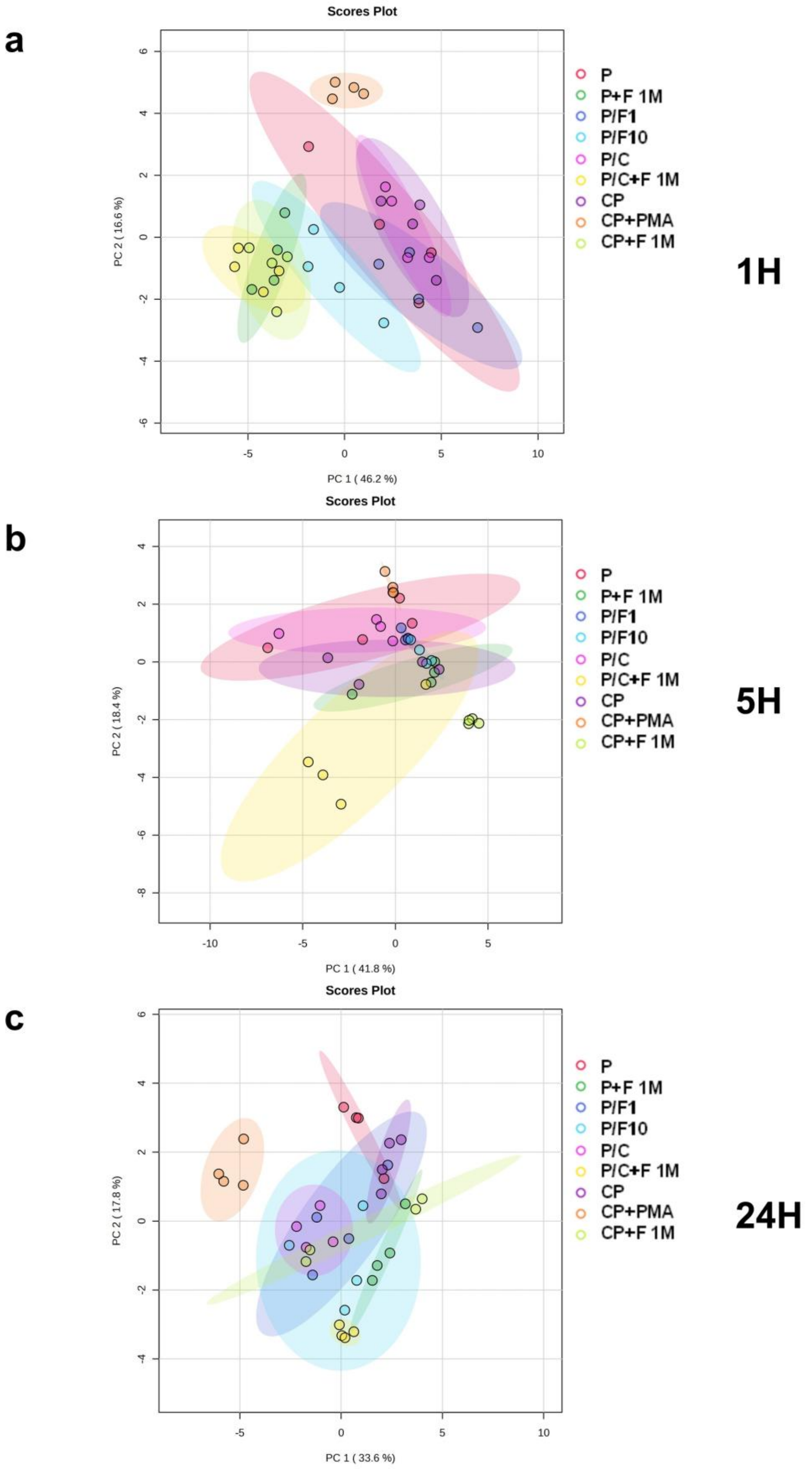


**Figure S3.** Principal component analysis (PCA) of the metabolomics data obtained at a) 1 hour, b) 5 hours, and c) 24 hours.

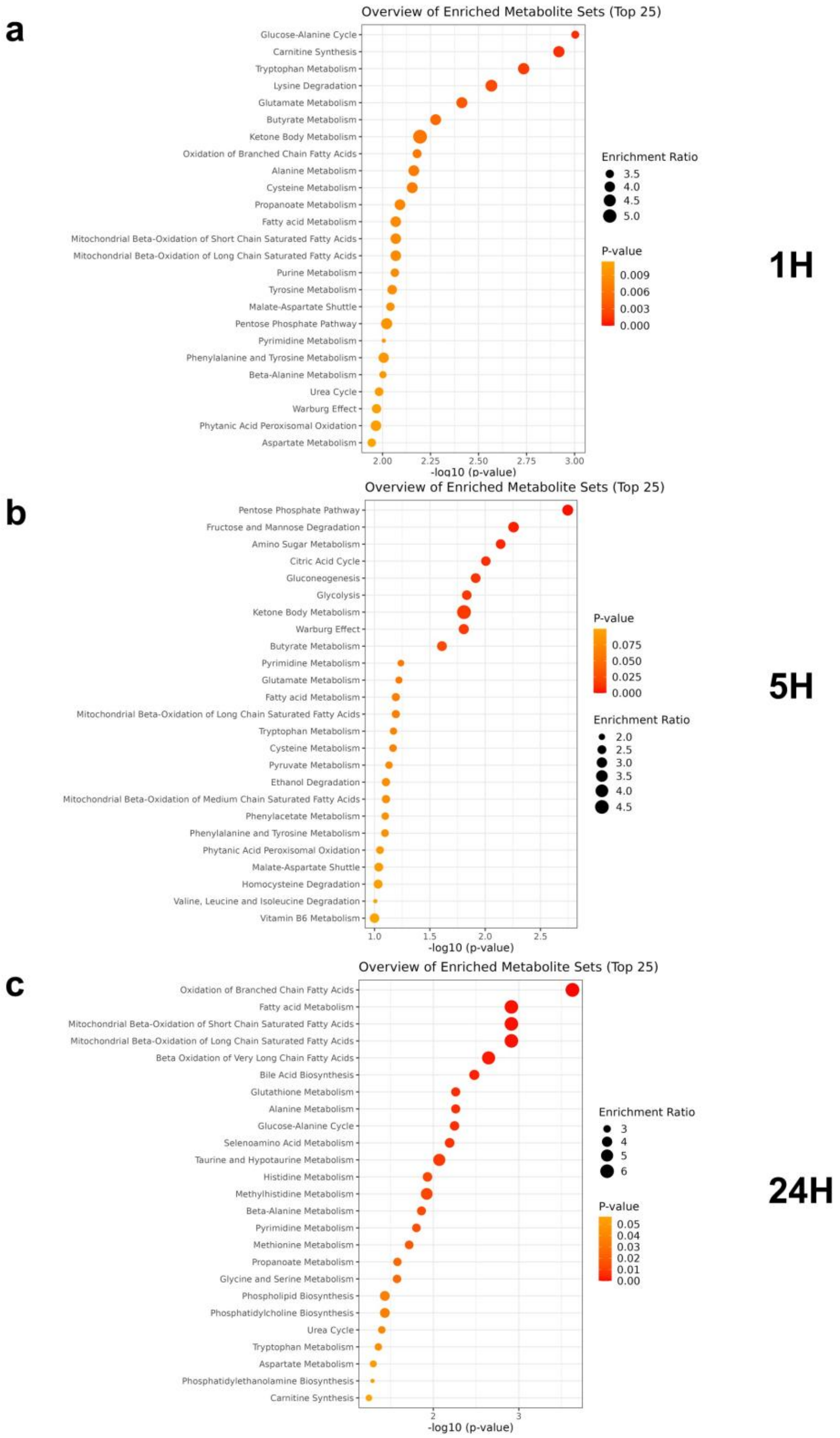


**Figure S4.** Pathway enrichment analysis of differentially expressed metabolites. Enrichment analysis of the differentially expressed metabolites obtained from the comparison of the experimental groups P vs P+F1M on the a) 1st hour, b) 5th hour, and c) 24th hour. The plot displays significantly enriched metabolic pathways based on over-representation analysis, ranked by pathway impact and statistical significance (–$\log_{10}$p-value). Pathways with $p < 0.05$ were considered significantly enriched.

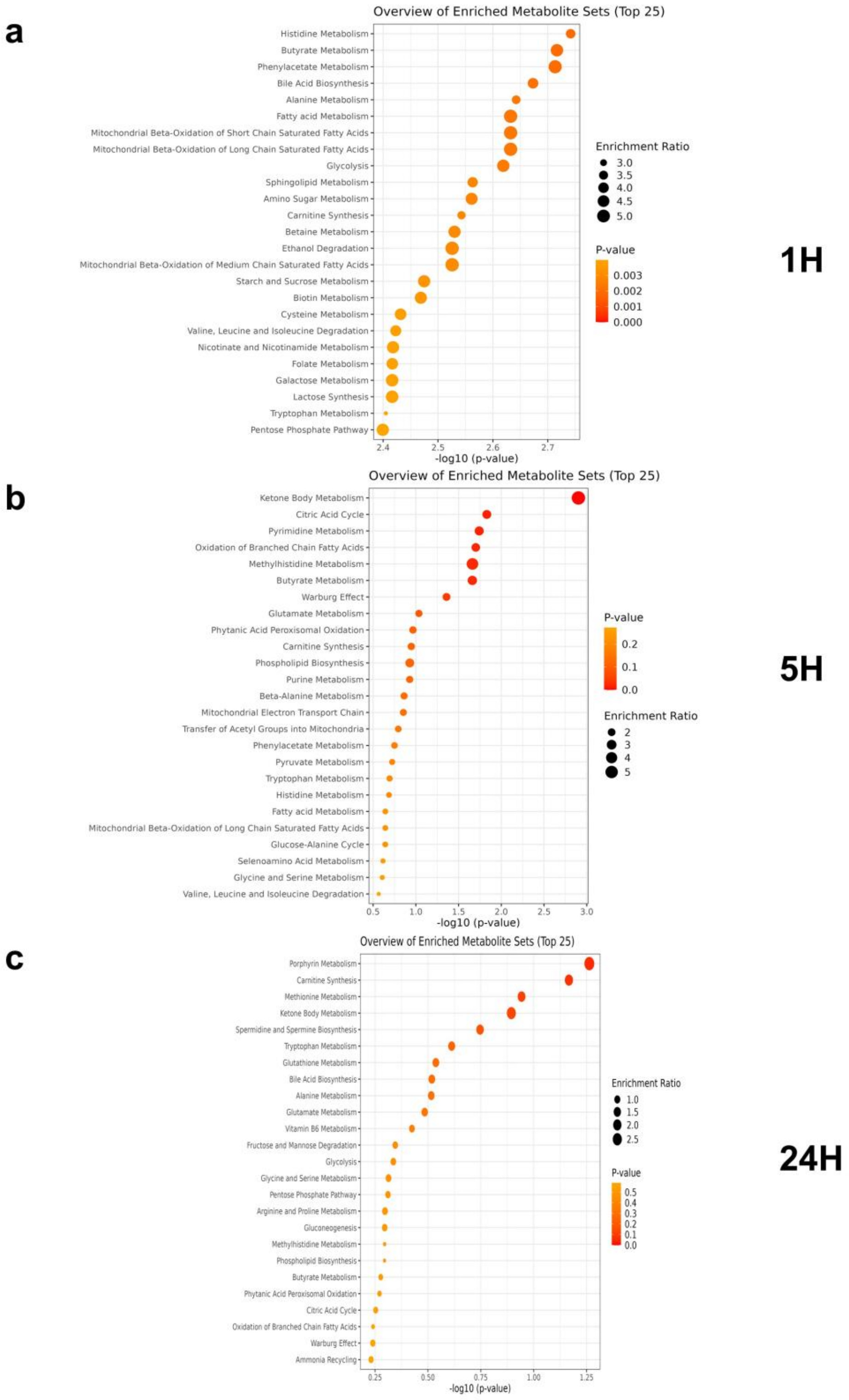


**Figure S5.** Pathway enrichment analysis of differentially expressed metabolites. Enrichment analysis of the differentially expressed metabolites obtained from the comparison of the experimental groups PCL vs FBS coated PCL on the a) 1st hour, b) 5th hour, and c) 24th hour. The plot displays significantly enriched metabolic pathways based on over-representation analysis, ranked by pathway impact and statistical significance ($–\log_{10}$p-value). Pathways with $p < 0.05$ were considered significantly enriched.

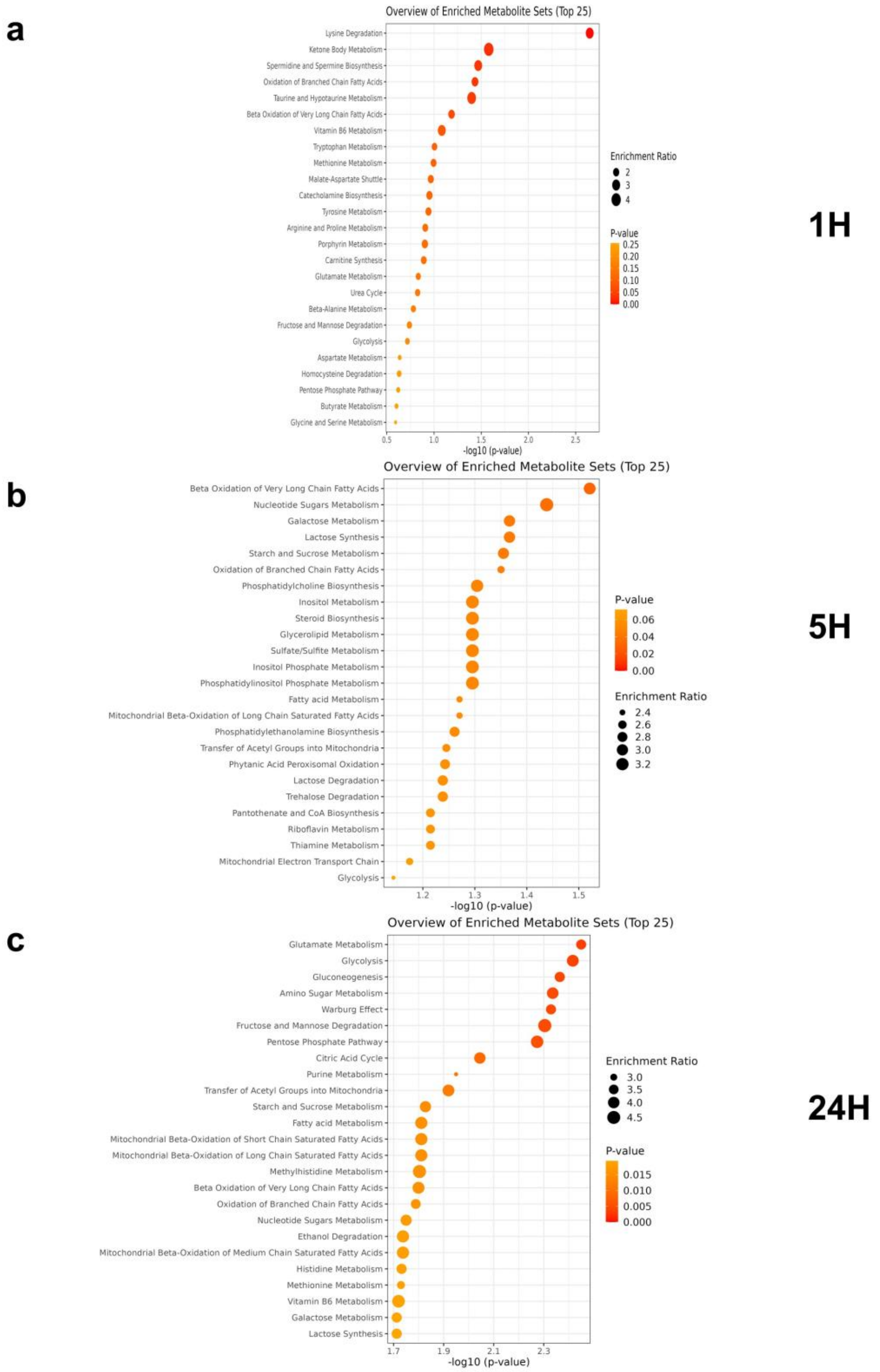


**Figure S6.** Pathway enrichment analysis of differentially expressed metabolites. Enrichment analysis of the differentially expressed metabolites obtained from the comparison of the experimental groups PCL vs collagen coated PCL on the a) 1st hour, b) 5th hour, and c) 24th hour. The plot displays significantly enriched metabolic pathways based on over-representation analysis, ranked by pathway impact and statistical significance ($–\log_{10}$p-value). Pathways with $p < 0.05$ were considered significantly enriched.

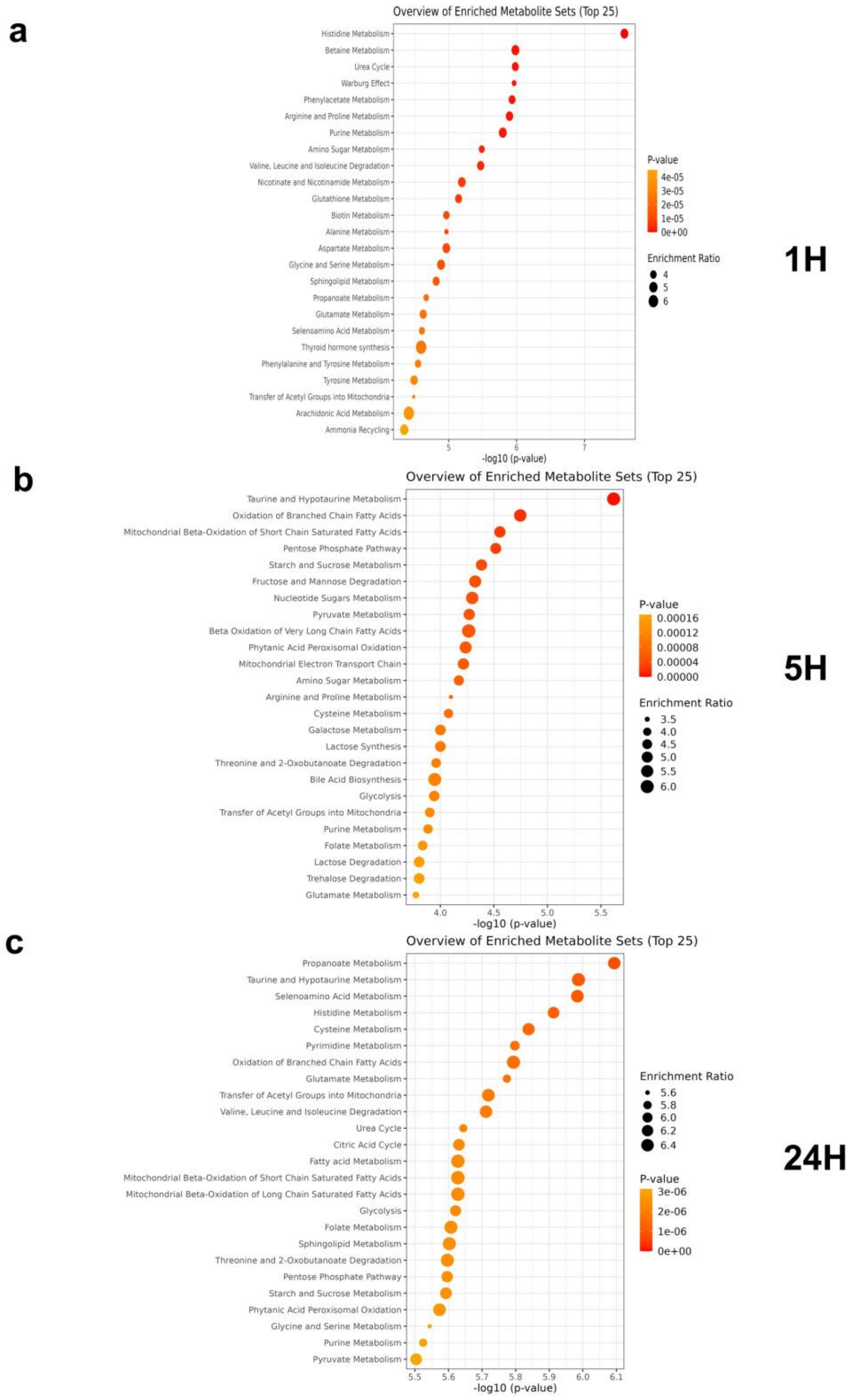


**Figure S7.** Pathway enrichment analysis of differentially expressed metabolites. Enrichment analysis of the differentially expressed metabolites obtained from the comparison of the experimental groups CP vs CP+PMA on the a) 1st hour, b) 5th hour, and c) 24th hour. The plot displays significantly enriched metabolic pathways based on over-representation analysis, ranked by pathway impact and statistical significance ($-\log_{10}$p-value). Pathways with $p < 0.05$ were considered significantly enriched.

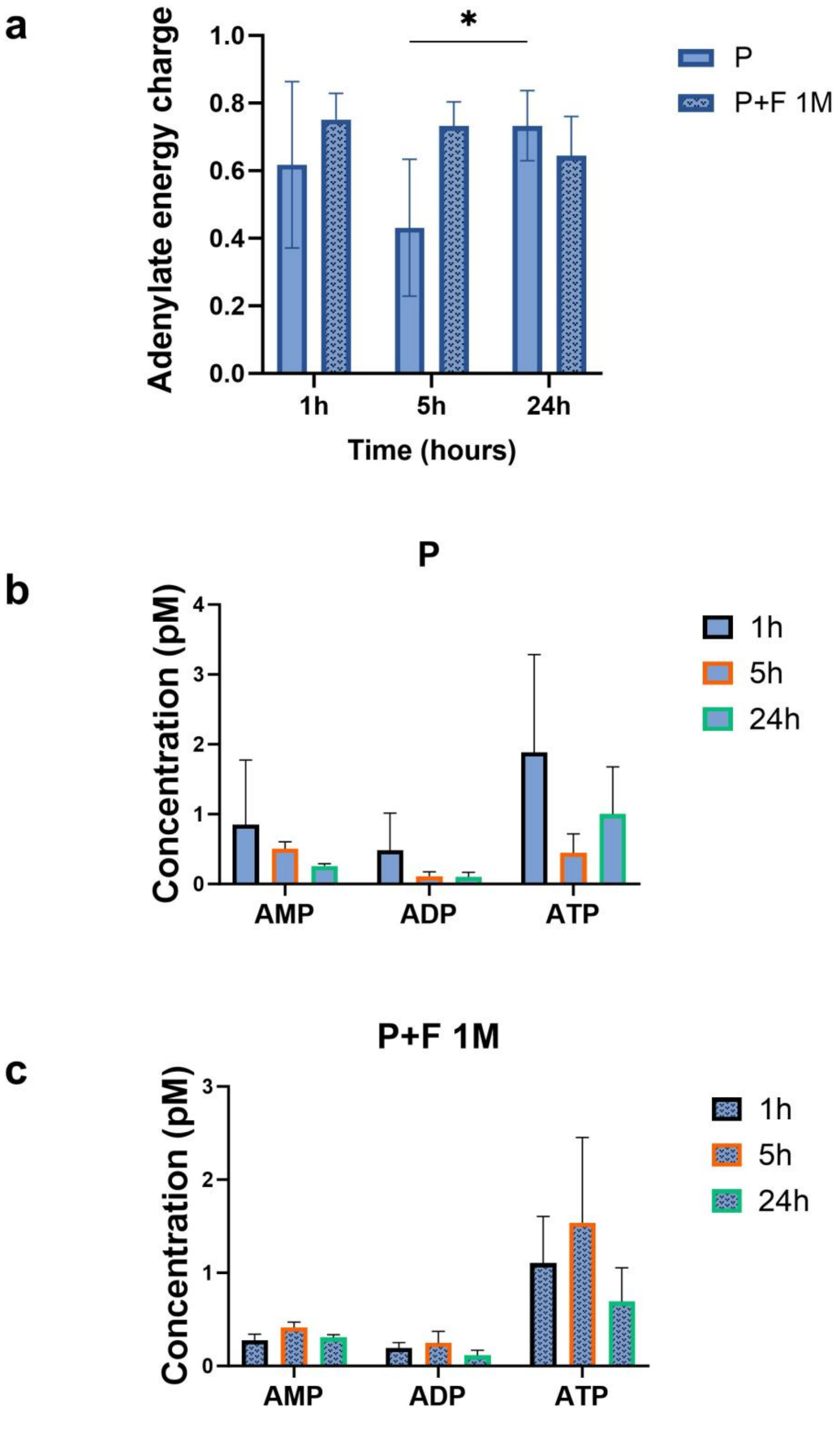


**Figure S8.** Adenylate metabolite dynamics in response to FBS. a) Adenylate energy charge (AEC, a metabolic indicator of intracellular energy status) in groups P and P+F 1M. b) Concentrations of adenylates (AMP, ADP, and ATP) at 1, 5, and 24 hours in neutrophils incubated on P. c) Concentrations of adenylates (AMP, ADP, and ATP) at 1, 5, and 24 hours in neutrophils incubated with P+F 1M. Column bars represent mean values +SD (SD: *: P< 0.05)

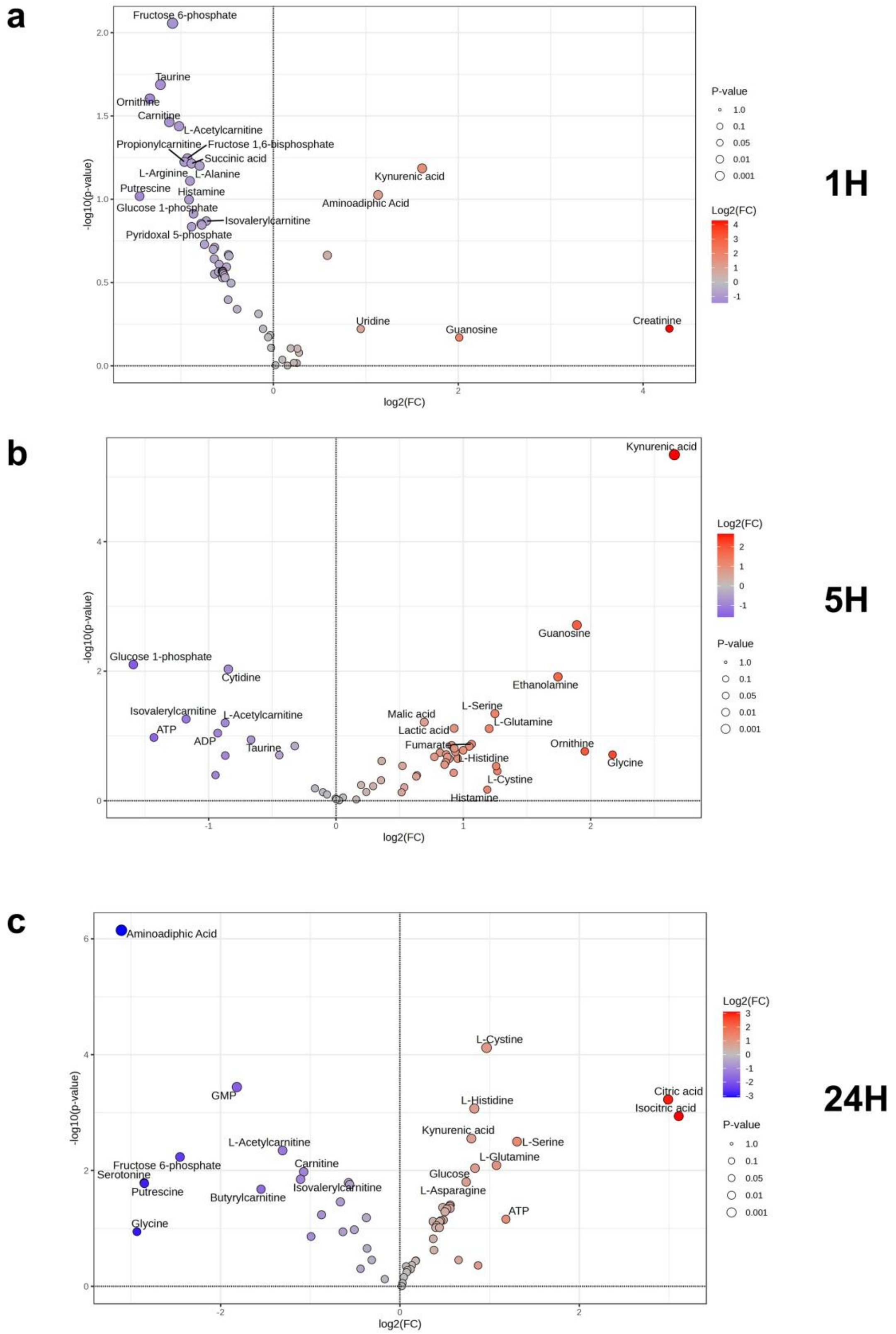


**Figure S9.** Volcano plots depicting differential expression of intracellular metabolites in neutrophils in response to PCL scaffold treatment alone on a) 1st hour, b) 5th hour, and c)

24th hour, identifying significantly altered metabolites based on fold change and adjusted p-values.

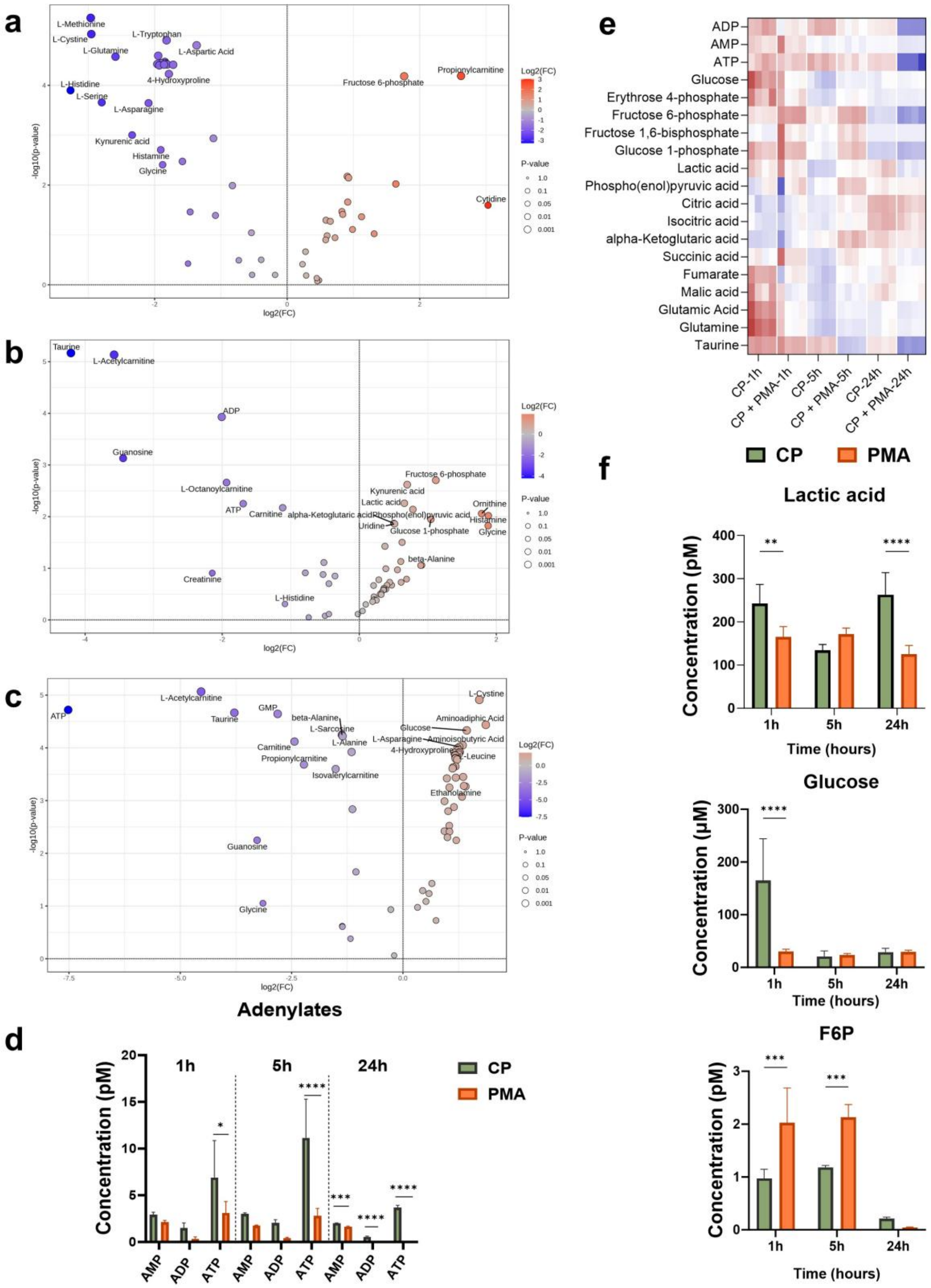


**Figure S10.** Metabolomic profiles of neutrophils following PMA treatment at defined time points. Volcano plots depict significantly upregulated and downregulated metabolites based on fold change and statistical significance at s) 1 hour, b) 5 hours, and c) 24 hours. d) Changes in intracellular adenylates (AMP, ADP, and ATP) from 1 to 24 hours. e) Heatmap illustrating the impact of PMA stimulation on neutrophil energy metabolism. f) Temporal comparison of lactic acid, glucose and fructose-6-phosphate (F6P) levels in neutrophils cultured on CP versus PMA-treated groups. Column bars represent mean values +SD (SD: **:P < 0.01, ***:P < 0.001, ****:P < 0.0001)

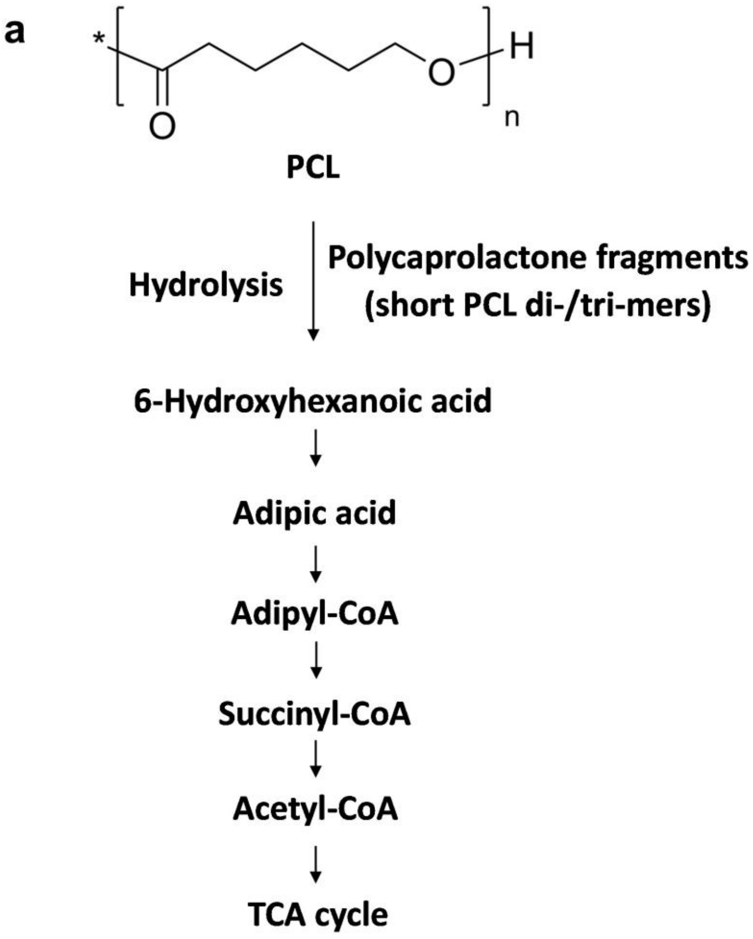


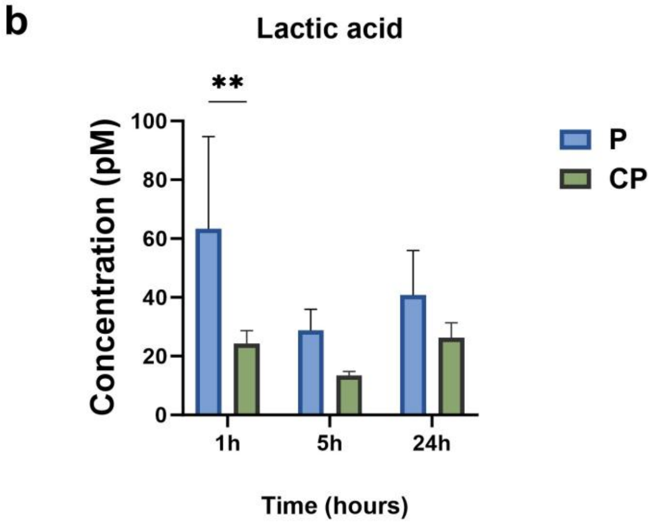

**Figure S11.** a) Schematic representation of polycaprolactone (PCL (P)) metabolization (The flowchart illustrates degradation processes involved in PCL breakdown), and b) Lactic acid concentration in neutrophils treated with P and CP. Column bars represent mean values +SD (SD: **: $P < 0.01$).
